# A Maximum-Likelihood Method to Construct Phylogenetic Trees Using Low-Homoplasy Markers: New Insights into Neoaves Phylogeny

**DOI:** 10.1101/2025.07.10.664098

**Authors:** Liang Gao, Stefan Grünewald

## Abstract

We present a novel probabilistic method for species tree reconstruction from low-homoplasy markers, termed MLRE (Maximum Likelihood for Retrotransposed Elements). MLRE uses an infinite-sites model for introducing new markers, which then evolve following a Markov process with three states: polymorphic, fixed, and lost. The model allows MLRE to optimize the likelihood of a tree with edge lengths in coalescent units. The method further incorporates variable parameters for insertion rates across lineages and missing data rates across taxa. For simulated data, MLRE consistently performs better than existing tree inference methods. Reanalyzing a retrotransposon dataset for Neoaves, we find better agreement with published trees from other data types, and we propose one strongly supported novel clade. These results underscore the value of retrotransposon markers in phylogenetics and highlight MLRE as a robust framework for species tree inference under challenging evolutionary conditions.

## Introduction

The problem to reconstruct the phylogenetic tree of Neoaves has garnered significant attention due to the extremely rapid radiation of bird lineages around the Cretaceous-Palaeoproterozoic extinction event [Claramunt and Cracraft, 2015], leading to conflicting phylogenetic results across studies. Jarvis et al. [2014] first provided a large-scale full genome analysis resulting in a neoaves tree with 48 taxa, followed by Prum et al. [2015], who examined 198 taxa. Disappointingly, the trees disagreed for most clades grouping together several orders. Since then, many new genomes have been sequenced, and new phylogenies have been proposed. Recently, Stiller et al. [2024] and Wu et al. [2024] prominently published phylogenies with 363 and 124 species, respectively, and they continue to reveal conflicting relationships within Neoaves. The differences are driven by data types, taxa sampling, and reconstruction methods [Reddy et al., 2017]. Despite all the conflict, several clades like landbirds (Telluvares, 13 orders) and waterbirds (Aequornithes, 7 orders) are consistently recovered throughout most or all studies. Reddy et al. [2017] classified Neoaves into seven such clades, the “magnificent seven” and the remaining three “orphan orders”.

Retrotransposable elements, a data type that is believed to exhibit extremely little homoplasy [Doronina et al., 2018], still contain many conflicting presence/absence patterns. This means that, unsurprisingly for a fast radiation, incomplete lineage sorting(ILS) is frequent [Matzke et al., 2012]. Suh et al. [2015] compiled a high quality data set with 2,118 retrotransposon markers for the taxa of Jarvis et al. [2014]. They constructed an MPRE (Most Parsimonious REtrotransposon) tree, but later Suh [2016] proposed a hard 9-way polytomy at the root of Neoaves.

However, a hard polytomy is not necessary to explain the conflicting or unresolved reconstructed trees, and that hypothesis has been disputed, for example by Reddy et al. [2017]. In addition to the problems posed by real data, like homoplasy and ILS, wrong model assumptions underlying the applied methods may introduce a systematic bias.

Homoplasy (parallel or reverse mutations) is the cause of the long branch attraction (LBA) bias of maximum parsimony [Felsenstein, 1978]. However, using maximum likelihood does not always avoid LBA. The problem can occur for short sequences [Huelsenbeck, 1995], partitioned likelihood [Roch et al., 2018], heterogeneity across sites [Lartillot et al., 2007], and various other reasons [Susko and Roger, 2021]. One main problem of ILS is the anomaly zone [Degnan and Rosenberg, 2006], a subset of trees with edge lengths where the most frequently observed gene tree is not the correct species tree. In practice, ILS make likelihood-based reconstruction from concatenated alignments inconsistent [Roch and Steel, 2015]. In the recent simulation with recombination of quartet trees in Zhang et al. [2025], concatenated likelihood is shown to suffer from long branch repulsion, the opposite bias of LBA.

If ILS was the only problem of a data set, then the Multi-Species Coalescent (MSC) model [Hudson, 1990, Degnan and Rosenberg, 2009, Jiao et al., 2021] would fix it. Exploiting the fact that there is no anomaly zone for rooted triplets, summary and site-based methods like ASTRAL [Mirarab et al., 2014, Mirarab and Warnow, 2015] and SVD Quartets [Chifman and Kubatko, 2014] are consistent in the presence of ILS [Mirarab et al., 2021]. However, the short sequences of single loci, heterogeneity across sites, and other problems of homoplasy make it hard to generate reliable input for those methods.

Since homoplasy, especially in combination with ILS, appears to be the dominant driver of species tree reconstruction errors, retrotransposons, the above-mentioned data type with extremely little homoplasy, appear to be the way out. They have been useful to resolve difficult phylogenetic relations [Nikaido et al., 1999, Nishihara et al., 2009], but currently available data sets often contain too few markers to yield significant results. The Suh [2016] data set has more than 2000 markers, but most of them confirm easy clades within the magnificent seven. To extract most information from such data sets, the choice of a method is crucial. Parsimony-based methods like MPRE that were initially preferred do not consider ILS and turned out to be biased in simulations [Molloy et al., 2022]. Two ILS-aware methods, ASTRAL BP and SDPquartet [Springer et al., 2020] exist, and they are adjustments of ASTRAL [Mirarab et al., 2014, Mirarab and Warnow, 2015] and SVD Quartets [Chifman and Kubatko, 2014] to retroelement insertion data. While they are shown to be consistent under the MSC model, their performance in an analysis of Suh’s data set [Gatesy and Springer, 2022] remained unsatisfactory, as the tree has little agreement with results from other data types and is very unstable with respect to removing few markers.

In this study, we propose MLRE, a likelihood-based method for estimating species trees using Retrotransposed Elements(RE) markers. It uses simple and intuitive assumptions to compute the likelihood of a data set for a rooted phylogenetic tree with edge lengths. Given a tree topology, the edge lengths can be optimised, and a standard tree search is employed to find a maximum likelihood tree. Variants of our model can handle variable insertion rates across lineages and different portions of missing data across taxa. Likelihood values can be used to quantify the differences between tree topologies.

For simulated data, MLRE performs clearly better than ASTRAL BP, if it is generated by our model, and slightly better for MSC-simulated data sets from Molloy et al. [2022]. For Suh’s Neoaves data set, we reconstruct trees that widely agree with many analyses of different data types and are more robust with respect to minor data set alterations than ASTRAL BP. Comparing previous publications, we find that trees by Jarvis et al. [2014] and Stiller et al. [2024] are in much better agreement with the RE data than Prum et al. [2015] and Wu et al. [2024].

## Materials and Methods

In this study, we introduce the Maximum Likelihood for Retrotransposed Elements (MLRE) method for estimating species trees using RE markers. It uses a novel probabilistic model based on the Infinite-sites Model assumption. Specifically, the model calculates the maximum likelihood values for different phylogenetic tree topologies, utilizing retrotransposon markers as input data. To identify the optimal species tree, a heuristic search algorithm is employed.

### Model assumptions

For a retrotransponson marker and a lineage, we distinguish the three states present (coded as 1), absent (coded as 0), and polymorphic (coded as 01). Every marker is introduced into a lineage by a retrotransposition into a single individual. This means that the state switches from absent to polymorphic. Through genetic drift, polymorphic markers can be fixed or lost, changing from 01 to 1 or 0, respectively. Therefore, the state 0 is used for the time before the insertion as well as after a loss. Since the number of potential markers is huge, the Infinite-sites model [Kimura, 1969] can be assumed, so every marker is inserted exactly once. Further, the insertion of a marker is extremely unlikely to be exactly reversed. If it is uncertain whether the insertion is present or absent in the taxon, the character “?” is used to represent missing data. The input consists of a presence/absence matrix where a set of markers assigns one of the observable states to each taxon.

Given a species tree with some taxon set, where every edge represents a lineage, the evolution of the markers is modeled by a probabilistic process with the following rules:

1. When the insertion event has not yet occurred, only 0 → 0 is possible.
2. 0 → 01 or 0 → 1 represent the appearance of the insertion, and this can only occur once during the entire evolutionary process.
3. 01 → 0, 01 → 1, and 01 → 01 represent the loss, fixation, or persistence of polymorphism in the population, respectively.
4. A marker that has been fixed will never be lost; thus, 1 → 01 or 1 → 0 do not occur. Similarly, once a marker is lost, it will never come back, and subsequent transitions will always be 0 → 0.

When mapping these retrotransposon markers onto the phylogenetic tree, the leaf nodes correspond to the input data, and the internal node states are inferred through different combinations that satisfy the leaf node labels. These combinations represent different potential evolutionary paths. To achieve the requirements of transition between two evolutionary states, the parent nodes of any two leaf nodes must follow the rules in Table 1. Note that there are only characters 0, 1 or “?” for the observed markers (leaf node), and there can be characters 0, 01, 1 and “?” for the internal node.

**Table 1.** Labeling rules for parent nodes.

| Descendent Nodes | Parent Node |
| --- | --- |
| At least one 0 and no 1 | 0 or 01 |
| At least one 1 and no 0 | 1 or 01 |
| Both 0 and 1 | 01 |
| Only ? | ? |

For example, if the observed labeling is (1, 1, 1, 0), there are six possible assignments of states to a 4-taxa caterpillar tree of the form (((1,1),1),0), as shown in Figure 1.

**Fig. 1:**
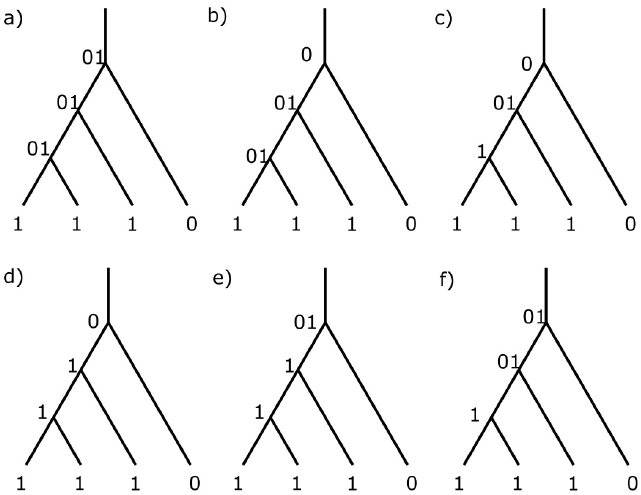
Six possible assignments of states to the interior nodes of an example tree and dataset.

For a given tree, all possible assignments of states to the interior vertices can be computed. Since the transition rules for evolutionary states at each node of the phylogenetic tree have already been outlined, the next step is to focus on the specific formulas that describe the probabilities associated with these transitions.

Under the infinite-sites assumption, insertions are irreversible, meaning each site can only undergo one insertion event. The probability of a marker to stay polymorphic is assumed to decay exponentially with time. The edge length for which this probability equals 1*/e* is defined to be one coalescent unit [Jiao et al., 2021], so for an edge of length *θ* the probability of staying polymorphic is *e*^−*θ*^. Over time, each marker either becomes fixed within the population or is lost.

For every lineage, we assume that the number of newly inserted markers is uniformly distributed, and the expected number equals the product of the edge length and an insertion rate *c*. As a unit of insertion rate, we use the number of markers that is expected to be polymorphic at the root. For the simplest version of our model, we assume an insertion rate *c* = 1 throughout the tree. Hence, there is an equilibrium number of polymorphic markers, and the expected number of fixed or lost markers during some period of time equals the expected number of newly introduced markers.

In order to compute transition probability matrices, we distinguish insertion edges from descendant edges for which both nodes are descendants of the introduction. For an insertion edge, the parent state is 0. The probability that a marker stays polymorphic depends on the portion of that edge that remains after the introduction. The expected number of new insertions appearing and maintaining polymorphism (0→01) over *θ* Is

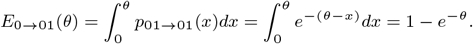

Here, *θ* represents the edge length, and *p*_01→01_(*x*) is the probability for a marker occurring at point *x* to maintain polymorphism for the remaining *θ* − *x*.

The total expected number of insertion events on an edge equals the product of the insertion and the edge length, which is *θ* for *c* = 1. Thus, the expected values for the other two outcomes — a marker occurring and being fixed (0→1) or lost (0→0) — are:

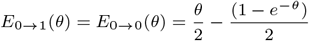

For a descendant edge, the probabilities of transitioning between the three evolutionary states are shown in Table 2. Specifically, the probability that the marker remains polymorphic through an edge of length *θ*, equals *e*^−*θ*^. The remaining probability, 1 − *e*^−*θ*^, is then equally divided between the two alternative outcomes: fixation and loss, each assigned a probability of 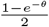. Realistically, the probabilities of fixation and loss may vary among different markers depending on the percentage of the population that carry the marker. For simplicity, we assume a symmetric model where the probabilities of fixation and loss are equal.

**Table 2.**
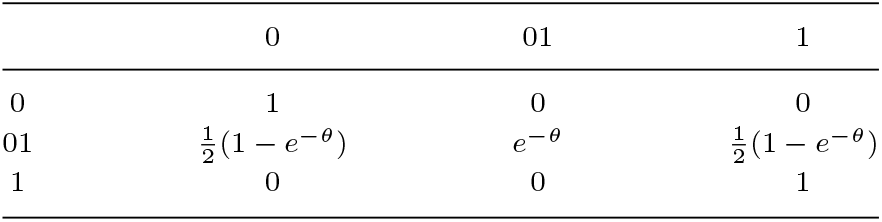
The transition probability matrix after insertion.

### Objective function

Based on the previous discussion, the expected marker occurrences and state transition probabilities for all edges in the evolutionary tree have been determined. A marker is an assignment of one of the states 0, 1,”?” to each taxon, and it is non-trivial if at least two taxa are assigned 1 and at least one taxon is assigned 0. Given a tree with interior edge length vector Θ = (*θ*_1_, *θ*_2_, …*θ*_*k*_), and a marker *m*, the expected number of occurrences of *m* can be computed by summing over all allowed labels of the interior vertices:

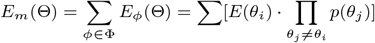

Here, *E*_*m*_(Θ) is the expected value of the marker *m*, Φ is the set of all allowed labels of the interior vertices, and the sum *E*_*φ*_(Θ) goes over all possible assignments in Φ that could lead to the observed marker pattern. For every *φ*, the marker is inserted on a specific edge *θ*_*i*_, with expected value *E*(*θ*_*i*_), and then transitions along other edges with probabilities *p*(*θ*).

In order to get the likelihood of the marker *m*, we have to divide the expected number of *m* by the sum of the expected numbers of all non-trivial markers. Then we multiply over all markers to get the likelihood of a data set:

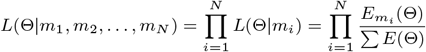

Here *{m*_1_, …, *m*_*N*_} is the set of all markers, ∑*E*(Θ) is the expected number of all non-trivial markers, and *L* refers to the likelihood of one or more markers.

### Optimization for Calculating Trivial Markers

As described in the objective function, we compute the expected value 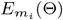 for each observed marker and sum over all non-trivial markers ∑*E*. For a species tree with *n* taxa, there are 2^*n*^ possible markers, among which only *n* + 2 are trivial. The remaining 2^*n*^ − *n* − 2 are non-trivial, and this number grows exponentially with *n*, while the number of trivial markers increases only linearly.

In practice, the number of observed markers in the input data is relatively small compared to the total number of non-trivial markers, with most non-trivial markers remaining unobserved. To avoid the computational burden of handling a large number of unobserved non-trivial markers, they can be replaced with trivial markers.

The expected number of all markers can be expressed as the sum of markers which are polymorphic at the root and markers that are introduced on internal edges of the phylogenetic tree. This sum equals

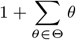

thus ∑*E* can be computed efficiently as

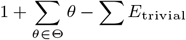

where ∑*E*_trivial_ is the expected number of all trivial markers. This approach to avoid computations for unobserved non-trivial site patterns has been used earlier to analyse morphological data [Lewis, 2001].

### Optional parameters

Our method comes with variants that allow grouping taxa together so that some leaves are polymorphic, variable insertion rates for different edges, and considering different missing data percentages for different taxa.

#### Group taxa and pendant edges

For large taxa sets, it is possible to replace stable clades by single taxa (see Preprocessing for details). In this case, such a group taxon might have 01 as its input state for a marker, if 0 and 1 both occur for different taxa within the group. More precisely, we define a consensus state for a taxon *s* representing a subset *S* of all taxa by:

1. If at least one taxon in *S* has state 1, and none has state 0, assign 1 to *s*.
2. If at least one taxon in *S* has state 0, and none has state 1, assign 0 to *s*.
3. If both state 0 and 1 occur within *S*, assign 01 to *s*.
4. If all taxa in *S* have state ?, assign ? to *s*.

Whenever 01 occurs as a state of a taxon *s* representing a group *S*, MLRE will compute an edge length for the pendant edge of *s*, and *s* represents the last common ancestor of *S*. The formulas for the expected values of the markers are refined correspondingly.

#### Insertion rate *c*

In reality, insertion rates can vary across different edges of the tree, depending on factors such as population size and population dynamics. For instance, in a rapidly growing population, markers are more likely to remain polymorphic, whereas a bottleneck would have the opposite effect. To account for this variability, a new parameter, *c*, is introduced, where the insertion rate for each internal edge is represented by *c*.

When an insertion occurs on an edge, the expectation for that edge must be multiplied by the corresponding *c*:

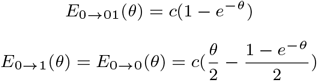

When *c* = 0, no markers are estimated to be inserted on this edge. When *c* → inf, then *θ* → 0, because the number of observed markers is finite. If short edges with very high *c* occur, then the model estimates that no previously polymorphic markers have been fixed or lost, but new markers have been inserted.

It is important to note that when using the insertion rate c, and simplifying the calculation with trivial markers, the expected number of markers should be modified as

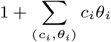

#### Missing-data rate *q*

Missing characters in markers are a fairly common problem in the real data. Their percentage varies between taxa, which may lead to bias.

To address this problem, we introduce one new variable *q* per taxon, where *q*_*i*_ represents the probability of a missing character in a taxon *i*. For each marker, the corresponding probability is adjusted by the character missing rate for each taxon. Thus, the original marker formula *E* is replaced by:

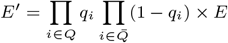

Here, *Q* is the set of taxa with state “?”, and 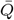 contains all other taxa.

Note that if the parameter *q* is used, it is not possible to use trivial markers to simplify the computation, since the number of trivial markers is then greatly increased.

### Pipeline and Implementation

#### Preprocessing

MLRE uses NEXUS as the input file format. In the preprocessing step, it identifies conflict-free subsets of taxa and optionally decomposes the tree reconstruction problem into several smaller instances. The output consists of one or more marker-tables.

In order to avoid infinitely long edges, maximal subsets of conflict-tree taxa are merged to form a new taxon. More precisely, two taxa are conflict-free if there is no marker where one taxon has state 1 and the other one has state 0. For every maximal conflict-free taxa set *S*, we replace *S* by a single new taxon *s* assigning consensus states to the markers as described in the previous subsection.

The next step of preprocessing makes MLRE suitable for large taxa sets. We use Buneman clustering to construct a (possibly highly unresolved) consensus tree. Then we produce separate datasets for every polytomy (a node with out-degree at least three) of the consensus tree. Every such dataset can then be resolved into a binary tree by the later steps of the pipeline, and these trees define a binary tree on the full taxa set.

A Buneman cluster is a subset of taxa, that is unanimously supported by all relevant triplets. Let #(*xy*|*z*) represent the counts of markers that are present in *x* and *y* and absent in *z*. If we have #(*xy*|*z*) *>* max*{*#(*xz*|*y*), #(*yz*|*x*)*}* for every choice of *x, y* ∈ *A*, and *z* a taxon outside *A*, then *A* is called a Buneman cluster. It is a straightforward and well-known consequence of earlier work, that all Buneman clusters can be computed efficiently, and they define a unique tree [Bandelt and Dress, 1986, Berry and Gascuel, 1997, Berry and Bryant, 1999].

We use binomial tests to define confidence values for every Buneman cluster *A*. For every choice of *x, y* ∈ *A*, and *z* a taxon outside *A*, let *k* = #(*xy*|*z*), and *n* = #(*xy*|*z*) + max*{*#(*xz*|*y*), #(*yz*|*x*)*}*. The triplet support of *xy*|*z* is the probability to observe less than *k* successes in *n* trials for a binomial test with success probability 0.5. The confidence value of *A* is minimum of all triplet supports obtained in this way. In MLRE, the user can set a threshold (default: 0.95), and the consensus tree contains precisely all Buneman clusters with confidence value above this threshold.

#### Searching for the Optimal Tree: NNI and SPR

The main part of MLRE searches for a binary tree with edge lengths, for an input dataset without conflict-free pairs of taxa. It involves computing a heuristic tree initially, then optimizing edge lengths and other parameters, and searching the space of tree topologies with tree rearrangement operations [Allen and Steel, 2001].

We first make a consensus tree of trusted clusters which will usually not be challenged during the tree search. By default, this is the previously mentioned Buneman tree with threshold 0.95. However, the user can change the threshold or input their own tree. We then use the triplet-joining method to fully resolve the constraint tree. Triplet-joining is an agglomerative heuristic similar to the Neighbor-Joining algorithm [Saitou and Nei, 1987]. It is a rooted version of the quartet-joining method, which was initially proposed by Berry [1997], and independently discovered several times [Ma et al., 2007], see Grünewald et al. [2009] for a consistency proof. Again, the user can specify a different initial tree. If such a tree conflicts with constraint tree, then the latter is replaced by the strict consensus of the user tree and the original consensus tree.

Once the initial tree is determined, the next step is to search for a better tree with a higher log-likelihood value. Given a current optimal tree, we use the Nearest Neighbor Interchange (NNI) operation to evaluate adjacent trees. As soon as a better scoring tree has found, it replaces the current optimal tree. Once an NNI-optimal tree has been found, we use the Subtree Pruning and Regrafting (SPR) operation. Since the size of the SPR neighborhood grows quadratically with the number of taxa [Song, 2003]. We only evaluate trees which contain all clusters of the consensus tree. As usual, an SPR-optimal tree is not guaranteed to be the global optimum.

#### Evaluation of Branch Support Using aBayes

Following the tree search, the resulting trees are further analyzed to assess the support for individual branches using aBayes [Anisimova et al., 2011]. Let *T*_1_ be the tree under consideration and *A* be a cluster of *T*_1_, and *T*_2_ and *T*_3_ be the two NNI neighbors of *T*_1_ that do not contain *A*. The branch support for *A* is then computed as:

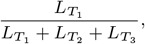

where 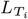 represents the likelihood of one of the three NNI trees.

##### Simulated Datasets

We provide a script (https://github.com/gaoliangs/MLRE) that simulates markers under the assumptions of our model using forward simulation. This simulation requires a given tree topology, branch lengths, and the parameter *c*. It outputs a CSV file containing the simulated marker data.

##### Implementation and Runtime Performance

The MLRE method is primarily implemented in Python, and MATLAB is used for likelihood maximization for each tree. The program can be downloaded from https://github.com/gaoliangs/MLRE.

The current implementation of MLRE is feasible for all available retrotransposon marker datasets. For the runtimes reported below, we use a standard laptop (MacBook Air M1).

For the 33-taxa Neoaves tree, MLRE started with an initial tree derived from the Buneman tree, fully resolved using the triplet-joining method. The search process involved 19 rounds of NNI moves, evaluating a total of 511 trees. The entire process took 3.5 hours to complete.

For the 33-taxa Neoaves tree with parameter *c*, the initial tree was also derived from the constraint tree and resolved using the triplet-joining method. The search included 22 NNI moves, examining 523 trees in total, and completed within 3.5 hours.

In terms of scalability, MLRE can handle large datasets depending on available memory. Our tests show that MLRE is capable of processing datasets with up to 500 taxa and 10,000 markers, demonstrating its feasibility for large-scale phylogenetic analyses.

## Results

### Simulations

#### Comparing the results of various tree-building methods

In this section, two quartet-based coalescent methods, ASTRAL BP and SDPquartet [Springer et al., 2020], are compared with the MLRE method. Both ASTRAL BP and SDPquartet are ILS-aware and utilize an RE matrix as input data.

For this comparison, a tree from real palaeognathae data was used as the basis for the simulation which served the same purpose in Molloy et al. [2022]. The difficulty of the reconstruction was increased by dividing all branch lengths longer than 0.1 by a factor of 10. We use the resulting tree once with constant insertion rate 1 and once with two edges with insertion rate 5 and half the original edge length and two edges with insertion rate 0.2 and twice the original edge length. These simulated true species trees with 12 taxa are illustrated in Figure S1. We then use our forward-time simulation to generate data sets with 100, 316, 1000, and 3162 markers. We use 100 replicate data sets for every choice of data size and fixed or variable *c*.

##### Accuracy of reconstruction

Figure 2 shows the average ratio of incorrectly reconstructed clusters. The results of ASTRAL BP and SDPquartet are extremely near to each other, and MLRE is significantly better than these two methods. As the data size increases, the error rates of all these methods converge to 0, and in Figure 2(b), the 3162 markers under MLRE with *c* gets 100% correct results. We conclude that 1) when data size rises, MLRE converges to the true tree. 2) MLRE is more accurate than the prior methods for all data sizes.

**Fig. 2:**
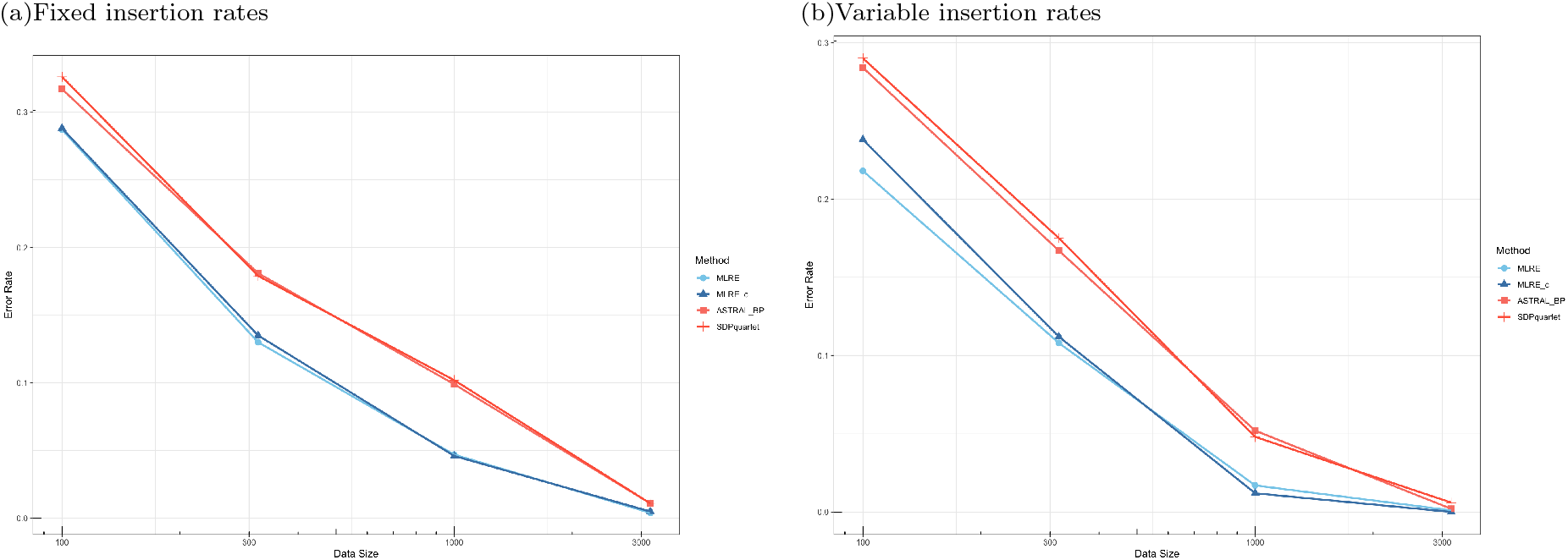
Comparison of three methods under the 12 taxa simulation tree. Error rates for MLRE and MLRE with and without variable *c* in blue, and ASTRAL BP and SDPquartet in red, data sizes are 100, 316, 1000 to 3162 markers.

In addition, we used the dataset from Molloy et al. [2022], which was simulated using the coalescent-based program ms [Hudson, 2002]. Figure S2 shows that the tree reconstructed using the MLRE method performs only slightly better than ASTRAL BP and most error rates are equal.

##### Branch Length Estimation

In addition to the accuracy of reconstruction, branch length estimates can also be compared. Since SDPquartet does not estimate branch lengths, we focus on comparing ASTRAL BP and MLRE. As shown in Figure 3, under identical conditions, MLRE consistently produces narrower box plots, indicating lower variance, and its median branch lengths are closer to the true values compared to ASTRAL BP(with one exception). Moreover, while MLRE estimates converge toward the true branch lengths with increasing data, ASTRAL BP exhibits systematic bias and tends to overestimate the edge lengths for large data sizes.

**Fig. 3:**
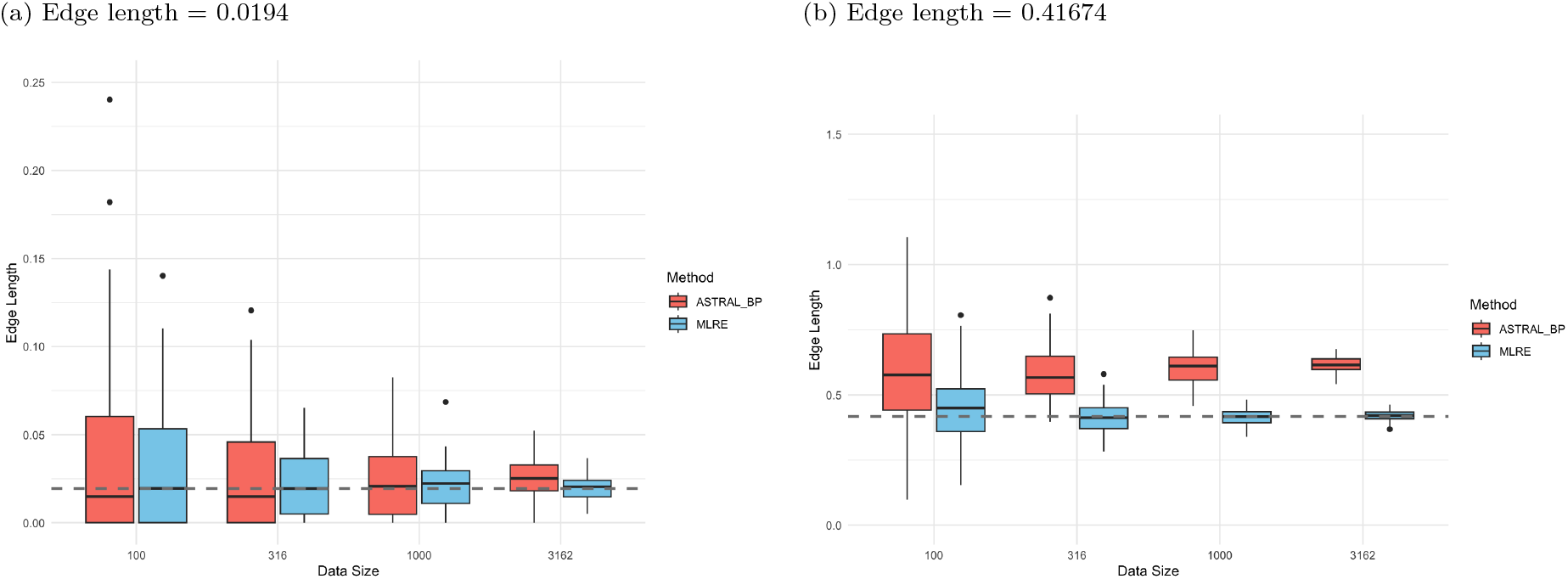
Distribution of Edge Lengths across Different Data Sizes and Methods. Results for edge length of (a) 0.019, and (b) 0.42. The blue boxplots represent MLRE estimates, and the red boxplots represent ASTRAL BP estimates. The gray horizontal line indicates the true branch length. The x-axis represents data size, and the y-axis represents the estimated edge length.

#### Data Size Required for Estimating the Insertion Rate *c*

MLRE shows higher accuracy than the other methods, with or without variable insertion rate *c*. However, simulations with variable *c* revealed that the estimated *c* values often deviated from the true values, particularly when the number of markers was small, leading to unstable estimates.

To assess the accuracy of *c*-estimation, simulations were conducted on a caterpillar tree ((((A,B),C),D),E) with two branch lengths (0.1 and 1). Three different values for *c* were tested (0.1, 1, and 10). For each scenario, datasets containing 100, 316, 1000, 3162, and 10,000 markers were generated to evaluate the performance of *c*-estimation across different tree sizes and branch lengths.

Figure S3 demonstrates that *c*-value estimates become increasingly accurate as marker numbers grow, although the impact varies with *c*-values and branch lengths. Larger deviations from *c* = 1, shorter branches, and smaller datasets lead to less stable and accurate estimates. However, increasing dataset size significantly improves stability, especially for longer branches.

#### The Impact of Missing Data

In this section, the impact of missing data (represented as question marks) on tree-building outcomes will be examined. The presence of question marks is common in real datasets, and their influence can complicate evolutionary analyses. For instance, in the neoaves dataset (Section Neoaves), the hummingbird has a notably high number of question marks (1,072 out of 2,118), while the penguins have less than 100.

To explore different missing data rates can miss lead tree reconstruction, a simulation was conducted using a 4-taxa tree with the following topology: (((A, B):0.1, C):1, D). In this simulation, taxon A has 50% missing data, while taxa B, C, and D each have 10% missing data.

In Figure 4, the results without the parameter *q* show that incorrect reconstructions are likely to occur. The error rate worsens as the dataset size increases. In contrast, when the parameter *q* is used, the accuracy of the reconstruction significantly improves, and the error rate converges to zero.

**Fig. 4:**
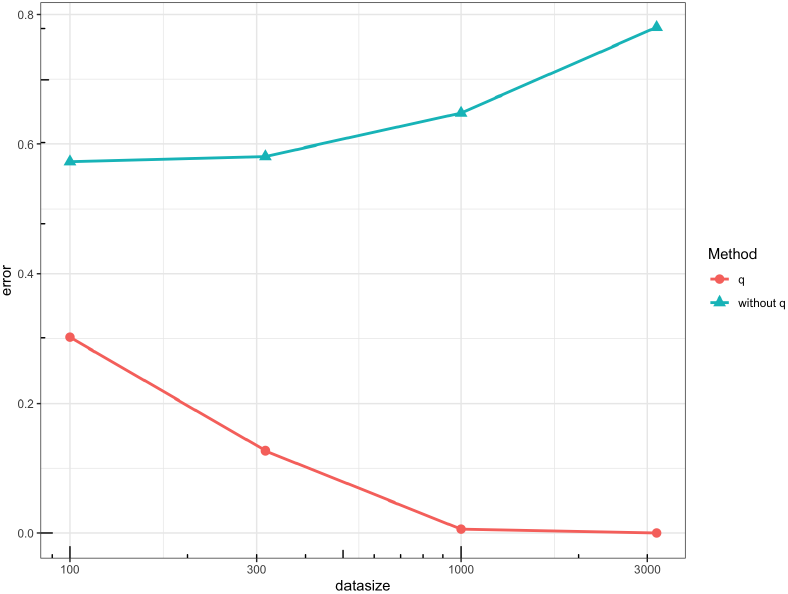
Error rate for reconstruction in the presence of missing data. The red line represents the use of the parameter *q* and the green line represents the nonuse of the parameter *q*. The x-axis represents the data size from 100, 316, and 1000 to 3,162 markers, and the y-axis represents the branching error rate of the reconstructed tree.

### Neoaves

#### Species Trees based on MLRE

MLRE is applied to the Neoaves data set originally collected by Suh et al. [2015], consisting of 2,118 retrotransposon markers from 48 species. Five of the 48 species are outgroups, containing only 0 or ‘?’ markers. Excluding these and merging the non-conflicting taxa, the dataset was reduced to 33 taxa, with the number of usable markers decreasing from the original 2,118 to 2,039. The Neoaves tree was then reconstructed using both constant *c* and variable *c* (Figure 5). Furthermore, when applying the Buneman cluster method (threshold: 0.95), we obtained 13 clades at the root, and the reconstructed tree is provided in Figure S15.

**Fig. 5:**
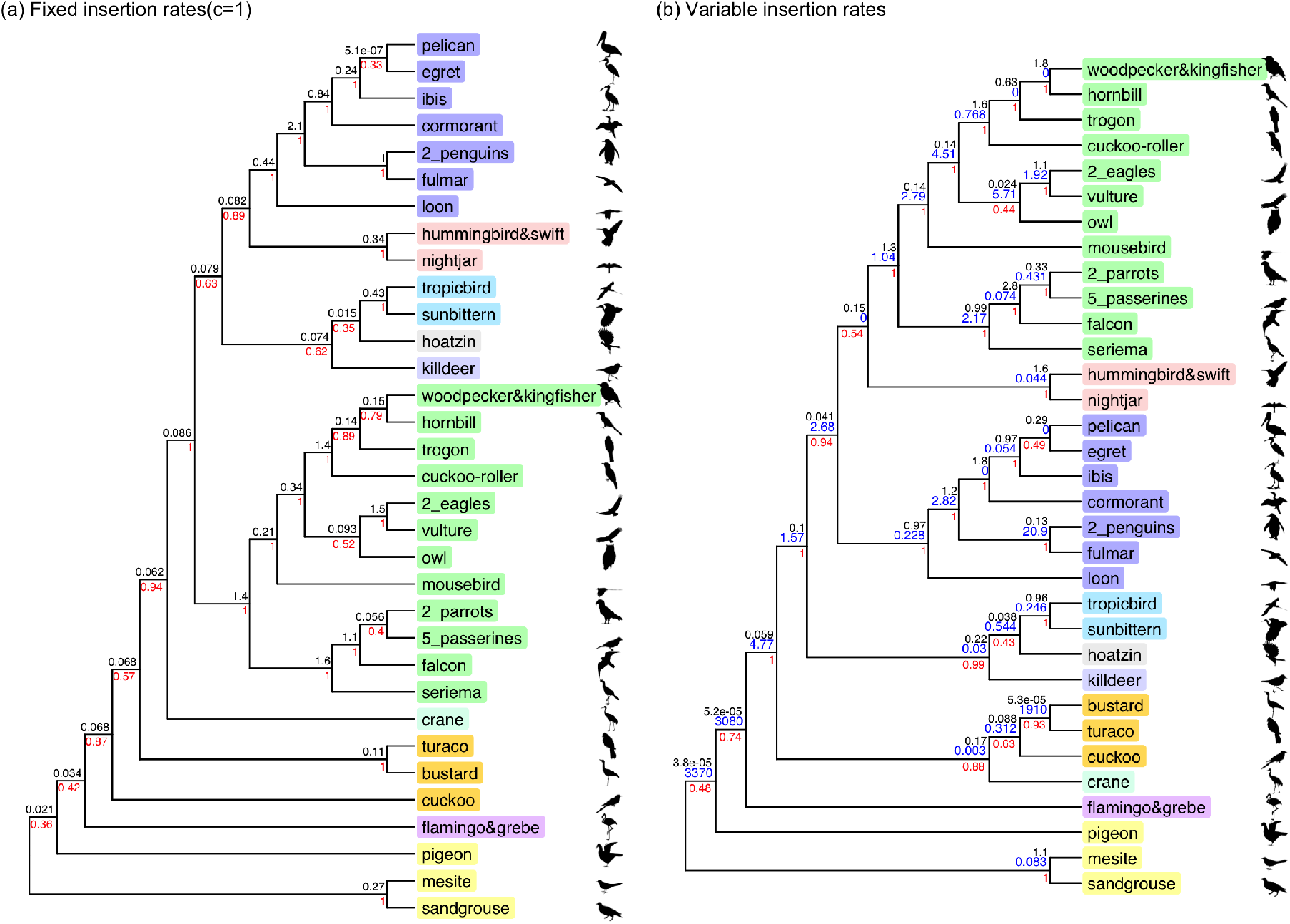
MLRE tree for 33 Neoaves taxa. (a) The Neoaves tree under the MLRE(c=1), logL = -10451.20647; (b) The Neoaves tree with variable insertion rates, logL = -9925.93296. Black numbers indicate branch length, blue numbers indicate insertion rates, and red numbers indicate branch support. The phylogenetic trees in this article are all plotted using the R package ggtree [Yu et al., 2017].

Near the root of Neoaves, Pteroclimesites consistently split first, but this branch is not well supported. The alternative topologies, namely Columbimorphae (Pteroclimesites + Columbiformes) or Columbiformes splitting first, both have logL-values very close to the optimal tree. Columbiformes and Mirandornithes are separated next. The clade of all other neoaves has been named Passerea, and it occurs in many previous studies [Jarvis et al., 2014, Reddy et al., 2017, Houde et al., 2019].

Another clade that is also common in other studies is Otidimorphae [Jarvis et al., 2014, Prum et al., 2015, Suh, 2016, Houde et al., 2019, Wu et al., 2024, Stiller et al., 2024], composed of Musophagiformes, Otidiformes, and Cuculiformes. In our study, Musophagiformes + Otidiformes is supported by 100% and 93% for constant *c* and variable *c*, respectively. However, Cuculiformes is only merged with the other two for variable *c*, and the support is around 63%. The insertion rate is estimated as 0.3 for Otidimorphae, and almost 2,000 for Musophagiformes + Otidiformes. This shape increasing might explain that Otidimorphae are not detected when *c* = 1 is assumed.

While the position of Gruiformes differ between the MLRE trees, the clade consisting of Aequornithes + Eurypqimorphae + Telluraves + Strisores + Charadriiformes + Opisthocomiformes is strongly supported by all variants. This combination has not been reported elsewhere, and it is the most confidently suggested novel clade of this study.

There is still a lot of uncertainty about the resolution within this big clade. Both trees in Figure 5 contain a cluster of Eurypqimorphae + Opisthocomiformes + Charadriiformes. It conflicts with frequently reconstructed Phaethoquornithes (Eurypqimorphae + Aequornithes), which is also detected by MLRE tree in Figure S15(a).

### Other Methods

#### MPRE tree from Suh et al. [2015]

Suh used Felsenstein’s polymorphism parsimony method [Felsenstein, 1979] to analyze the presence/absence matrix of retrotransposons (REs) to heuristically obtain an MPRE tree. The MPRE tree has 1,398 congruent markers (Figure 6(a)), and its tree length (a measure of total independent allele fixations, defined by Suh) is 5,377. For comparison, mapping these 2,118 markers to the Jarvis tree will yield 1,373 congruent markers and tree length = 5,579.

**Fig. 6:**
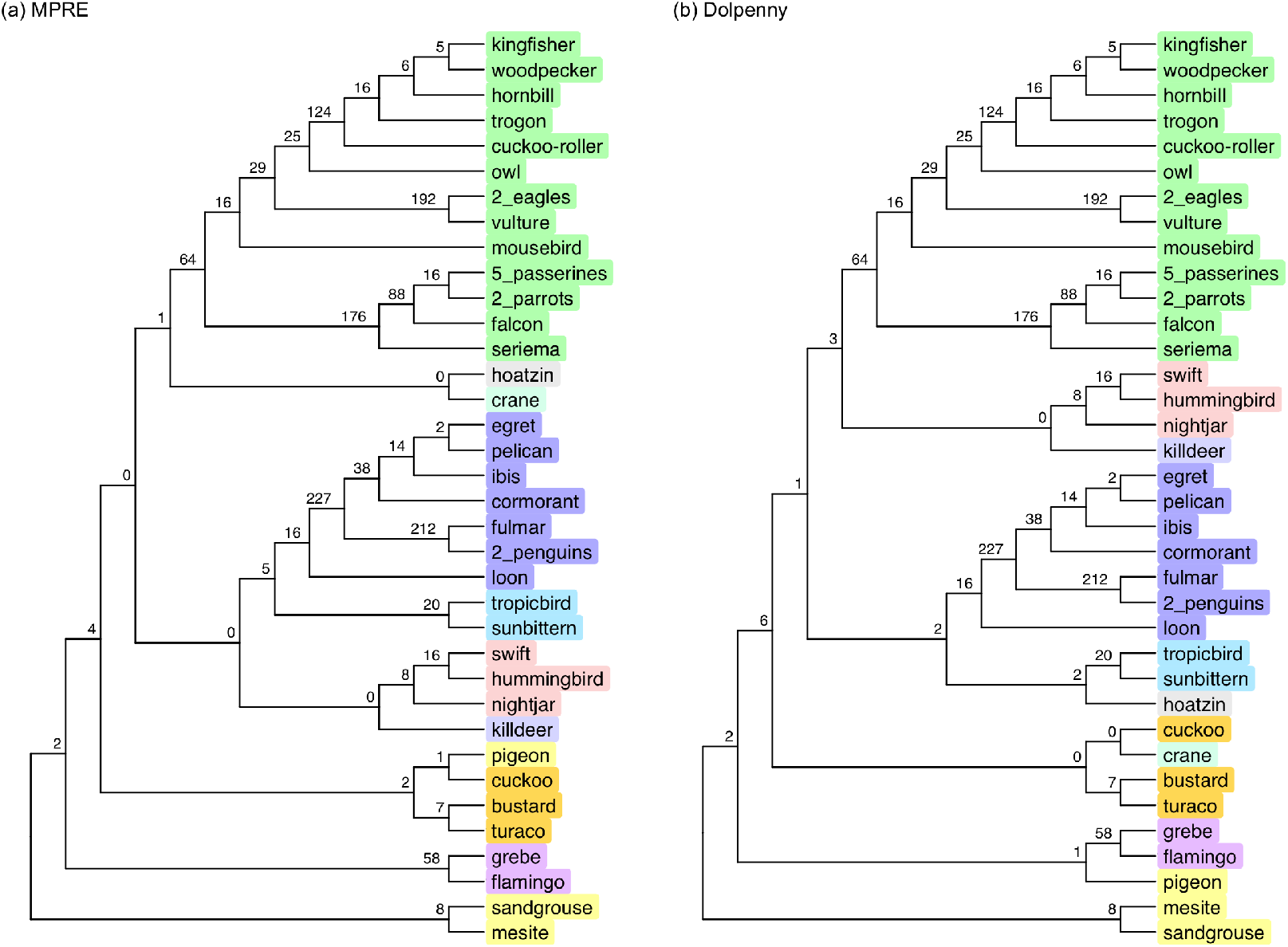
(a) MPRE tree; (b) Optimal tree under Polymorphism Parsimony(Dolpenny). Branches are labeled by the number of supporting markers. There are 1398 markers in the MPRE tree and 1400 markers in the best tree.

Suh did not find the optimal tree under this criterion. We used Dolpenny, an exact branch-and-bound version of polymorphism Silhouettes behind each species in the figure are from: http://phylopic.org/ parsimony in PHYLIP [Felsenstein, 1989], that can deal with small taxa numbers. By identifying turaco and bustard from the Buneman taxa (in Figure S15), we got a 12 taxa dataset. For conflicting markers in the merged taxa, polymorphic states were retained and denoted as P. We replaced group taxa by the subtrees from Suh. This new tree has 1,400 congruent markers (Figure 6(b)) and a tree length of 5,355.

While Suh’s tree includes the Phaethoquornithes clade, it is missing in the optimal tree. This suggests that the Phaethoquornithes clade only appeared in Suh’s results by chance. The MPRE method does not support other commonly recognized clades such as Otidimorphae, and Columbimorphae found in previous studies.

The new optimal tree includes clades such as Aequornithes + Eurypygimorphae + Telluraves + Strisores + Charadriiformes + Opisthocomiformes and Musophagiformes + Otidiformes + Cuculiformes + Gruiformes, which are more consistent with the outcomes of the MLRE method compared to the MPRE tree.

Additionally, the MLRE tree has 1,404 congruent markers (6 more than the MPRE tree) (Figure S14). When comparing the MLRE and MPRE methods, the score function of MLRE is more meaningful because it accounts for differences in branch lengths. For shorter branches, markers are more likely to remain polymorphic, while MPRE treats all edges equally, leading to bias.

##### ASTRAL BP & SDPquartets from Gatesy and Springer [2022]

ASTRAL BP calculates species trees by scoring bipartitions based on their frequency across multiple gene trees, using dynamic programming to efficiently find the optimal species tree while handling incomplete lineage sorting. SDPquartet uses quartet analysis to infer species trees by examining the relationships among sets of four species at a time, combining these quartets to construct a consistent global species tree while accounting for incomplete lineage sorting. Gatesy and Springer [2022] got the same tree with both methods.

However, when repeating the experiment using the same data and methods, we observed that out of the 2,118 markers, 817 markers have a “?” for all five outgroups, with no “0” present. For such markers, ASTRAL BP and SDPquartets miss the information that 0 is the ancestral state. We therefore added an additional outgroup with all markers assigned as 0.

After adjusting the input data, new ASTRAL BP and SDPquartet trees were obtained, respectively (Figure 7). The adjusted phylogenetic trees show some improvements under both methods. While originally Strisores was placed near the root, now the clade moved to the lower part of the tree, and Pteroclimesites became the clade closest to the root, which is more consistent with the common results in other phylogenetic analyses of Neoaves. As shown in Table 4, the log-likelihood difference between MLRE and Gatesy and Springer [2022] (Figure S10), which is approximately 70, decreases to about 30 between MLRE and adjusted ASTRAL BP (Figure S12).

**Fig. 7:**
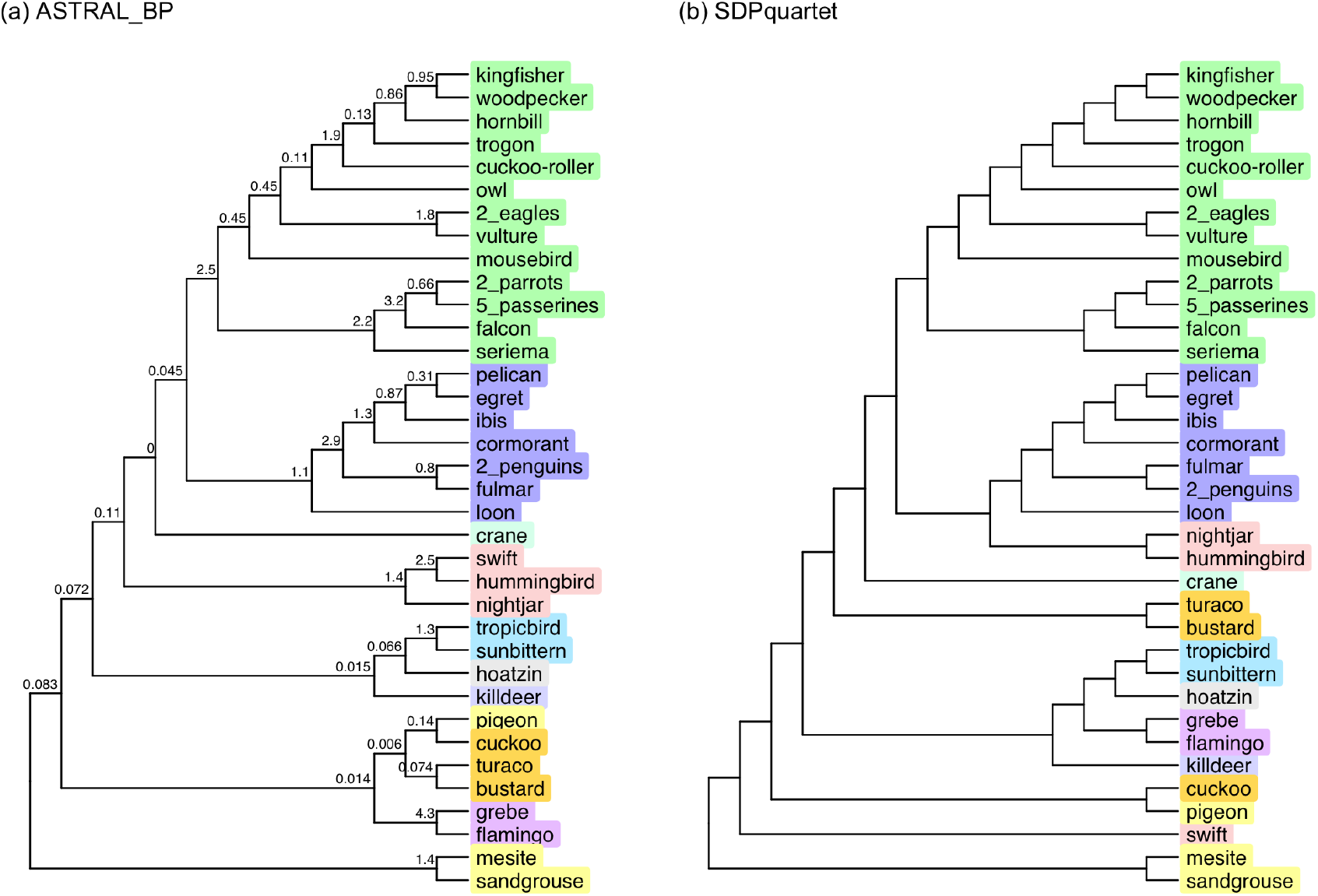
(a) Adjusted ASTRAL BP tree; (b) Adjusted SDPquartet tree. The branch length in CUs are estimated by ASTRAL. In the SDPquartet tree, the swift fails to be grouped with the two other species in Strisores, hummingbird and nightjar.

However, many commonly recognized clades such as Otidimorphae, Columbimorphae, and Phaethoquornithes were not successfully reconstructed. Additionally, the SDPquartet method revealed a clear mistake: it failed to correctly place the three species within the Strisores clade — hummingbird, nightjar, and swift—into the same evolutionary branch.

#### Stability

To evaluate the stability of the MLRE algorithm, several key markers were added or removed to determine whether the final results differed from the current ones.

##### Remove two critical markers

Gatesy and Springer [2022] highlighted two critical markers whose removal leads to the loss of many key evolutionary clades when analyzed with the ASTRAL BP method, resulting in a completely different phylogenetic tree.

- Marker 1 includes: hoatzin, sunbittern, tropicbird, grebe, flamingo, bustard, turaco, pigeon, and cuckoo.
- Marker 2 includes: mesite, hoatzin, sunbittern, tropicbird, grebe, flamingo, killdeer, cuckoo, and crane.

Upon removing these two markers, the analysis using ASTRAL BP revealed that although the Buneman clustering and its internal topologies remained stable, only 2 out of the 11 evolutionary clades generated from the 13 taxa at the root of the Neoaves phylogenetic tree were retained. This indicates that the ASTRAL BP method exhibits a certain degree of instability when handling the removal of critical markers.

In contrast, after removing the two markers, the optimal tree generated by MLRE with or without variable *c* remained consistent with those obtained before marker removal.

##### Add some markers supporting clusters from other studies

Several clades from previously published studies, including Jarvis et al. [2014], Prum et al. [2015], Stiller et al. [2024], and Wu et al. [2024], were selected for testing. We then added markers that are present precisely for the taxa in the cluster until the cluster is included in the MLRE. Table 3 presents the additional number of markers required to reconstruct these clades in the phylogenetic analysis. The specific taxa included in each cluster are listed in Supplementary Table S1.

**Table 3.** Number of markers added.

| Cluster | variable $c$ | constant $c$ |
| --- | --- | --- |
| Otididae | 3 | 3 |
| Otidimorphae | 0 | 1 |
| Columbea | 8 | 4 |
| Columbimorphae | 2 | 2 |
| Gruae | 3 | 4 |
| Phaethoquornithes | 1 | 1 |
| Inopinaves | 6 | 3 |
| Gruimorphae | 3 | 3 |
| Columbaves | 9 | 9 |
| Litusilvanae | 3 | 5 |
| Elementaves | 4 | >10 |
| Aequorlitorntithes | >10 | >10 |

Compared to previous studies, although some of our results do not include the three common clades, Otidimorphae, Columbimorphae, and Phaethoquornithes, our method is capable of generating an evolutionary tree that includes these clades by incorporating few markers that contain them. Specifically, our approach can partially predict these clades with the original data and can stably detect them after the addition of one or two markers. Thus, all three clades represent competitive alternative clusters, and their absence can be explained by the limited available data. All other clusters need at least 3 markers. Especially, the frequently discovered Columbaves is not supported by this data set.

### Score comparison with previous studies

We applied the MLRE method to evaluate several previously published Neoaves phylogenies, as well as our own adjusted ASTRAL BP and Dolpenny trees. As shown in Table 4, all these trees yield substantially lower MLRE scores compared to the optimal tree inferred by our method. Moreover, some branches have MLRE-estimated lengths equal to zero (Figure S4–S13), suggesting weak support under our model. We count edges if the edge length as well as the product of insertion rate and edge length are both less than 10^−4^.

**Table 4.** Trees from other studies.

|  | logL-value | length=0 edge |
| --- | --- | --- |
| Jarvis et al. [2014] | -9971.58 | 3 |
| Prum et al. [2015] | -10105.12 | 2 |
| Reddy et al. [2017] | -10015.38 | 4 |
| Kuhl et al. [2020] | -10019.76 | 2 |
| Stiller et al. [2024] | -9986.85 | 3 |
| Wu et al. [2024] | -10068.39 | 2 |
| Gatesy and Springer [2022] | -9979.91 | 1 |
| MPRE [Suh et al., 2015] | -9940.25 | 0 |
| Dolpenny | -9936.35 | 0 |
| ASTRAL_BP(adjusted) | -9947.14 | 0 |
| MLRE | -9925.93 | 0 |

Among the evaluated trees, those from Prum et al. [2015] and Wu et al. [2024] show the worst fit to the MLRE model. Generally, trees made from the same data set tend to have a better score than phylogenies from different data types. Among the latter, the best score is obtained by Jarvis et al. [2014] which uses the same taxa set.

### Landbird Problem

The landbirds contain the majority of all ling bird species, and the reconstruction of their phylogeny has driven particular interest. There are five well-established clades, Australaves (falcons, parrots, songbirds, and more), Cavitaves (woodpeckers, hornbills and more), Accipitrimorphae (eagles and New World vultures), Strigiformes (owls), and Coliiformes (mousebirds). Resolving the relation between these five taxa is difficult, and the two most widely discussed questions are the position of the mousebirds and whether owls are the sister group of Cavitaves or Accipitrimorphae.

#### The evidence clearly supports that mousebird is the sister taxon to the remaining Afroaves

Most publications since the arrival of full genomes have grouped mousebirds with Cavitaves, for example Jarvis et al. [2014], Prum et al. [2015], Stiller et al. [2024], and Wu et al. [2024], and the union of these clades has been defined as Coraciimorphae by the PhyloCode Sangster et al. [2022]. However, it was already pointed out by the original analysis of this data set Suh et al. [2015] that retrotransposons do not agree with the grouping. Instead, all publications analysing the data find that mousebirds are the sister taxon to the remaining Afroaves. The result of applying MLRE to address the mousebird question is clear. In Figure 8, (a) and (b), the lanndbird MLRE trees with constant and variable insertion rate *c* are shown. The topology is the same and the cluster of all other Afroaves than mousebird is strongly supported with 100% aBayes value.

**Fig. 8:**
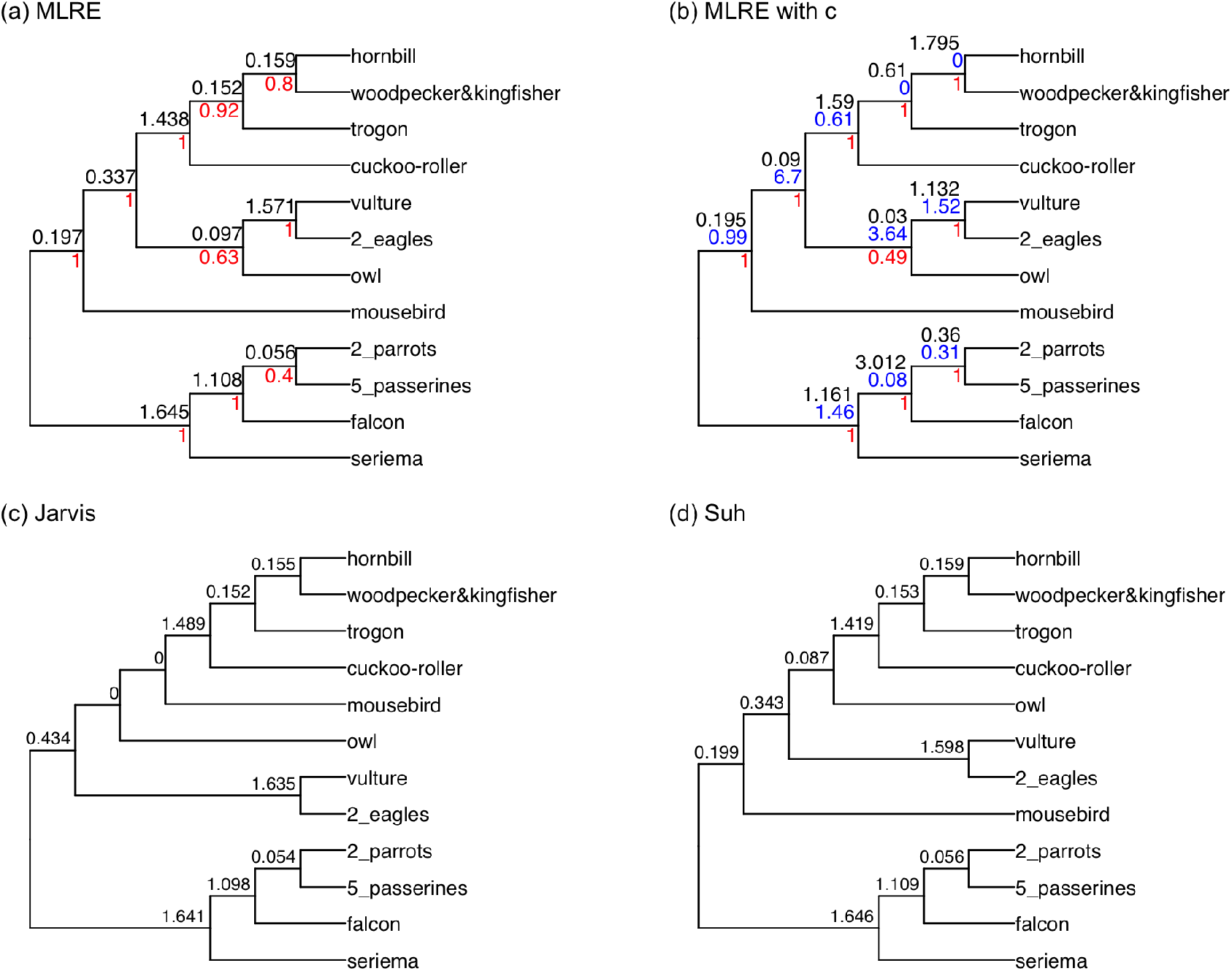
Trees for 12 landbird taxa. (a) MLRE; (b) MLRE with c; (c) Landbird tree extracted from the Jarvis et al. [2014] main tree; (d) Landbird tree from Suh et al. [2015] MPRE tree.

In addition, MLRE was applied to analyze the landbird tree from Jarvis et al. [2014]. The tree contains two edges conflicting the MLRE tree, and both receive zero branch length. The tree generated by MLRE achieved -3150.82826, and the Jarvis tree only reached -3187.53116.

#### The missing characters within landbirds do not affect the placement of mousebird

One potential problem that could mislead the placement of mousebirds in the landbird tree is the uneven distribution of missing data. Specifically, among the five taxa within landbirds, two independent taxa exhibit a relatively high proportion of missing characters: mousebird with 83/310 and owl with 34/310. In contrast, the composite taxa—Cavitaves, Accipitrimorphae, and Australaves—have significantly fewer missing characters, with 6/310, 2/310, and 2/310, respectively.

Even with the inclusion of parameter *q*, the results still indicate that the two conflicting branches of the Jarvis tree have zero branch lengths, and the logL-value (logL = -1491.50680) is substantially lower than the logL-value of the landbird tree generated by MLRE (logL = -1458.65812, Figure 9). This demonstrates that despite approximately 27% of characters being missing in mousebird, these missing characters have no substantial impact on the placement of mousebird in the phylogenetic tree.

**Fig. 9:**
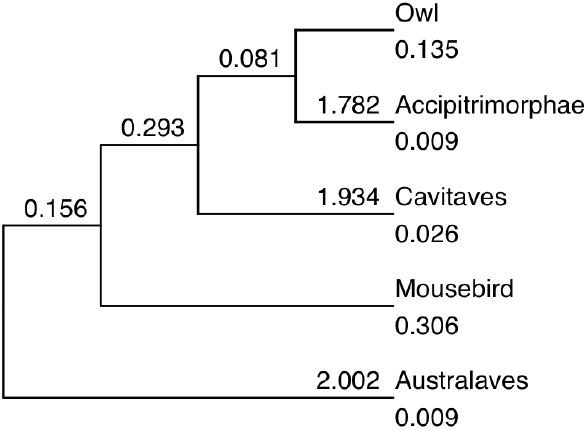
Landbird tree with variable missing data rates. Branches are labeled by the edge length. Missing data rates are displayed behind the taxa names.

#### The placement of owls: Cavitaves or Accipitrimorphae?

The placement of the owls within the landbirds is difficult, and two different positions have frequently been published. Owls are either grouped with Cavitaves [Jarvis et al., 2014, Prum et al., 2015] or with Accipitrimorphae [Houde et al., 2019, Stiller et al., 2024, Wu et al., 2024]. The third alternative, owls being the outgroup to a cluster of Cavitaves and Accipitrimorphae has not been found throughout different methods and data types. Suh et al. [2015] already found the support for the two conflicting hypotheses, and Stiller et al. [2024] observed the same for whole genome alignments, suggesting an ancestral hybridization.

While Suh’s MPRE tree prefers to group owls with Cavitaves (Figure 8d), MLRE chooses to put them with Accipitrimorphae. Both trees have similar edge lengths, and the log-likelihood differs by less than one. We used several choices of taxa sets, the 12 landbird taxa, Buneman clustering with a threshold of 0.99 to get a reduced taxa number of 5, and trees on all 33 taxa that only differ in the placement of owls, and we changed the optimized parameters.

Regardless of the presence of the insertion rate *c*, the combination of Accipitrimorphae and owl yielded slightly better scores compared to the combination of Cavitaves and owl when using the 12 landbird species alone as taxa. However, when the analysis was conducted using the five group taxa, the combination of Accipitrimorphae and owl produced better results when *c* = 1. In contrast, when *c* was treated as a variable parameter, the combination of Cavitaves and owl provided better results.

The results in Table 5 indicate that owl + Cavitaves shows a slight advantage only when using composite taxa and variable insertion rate *c*. In all other cases, owl + Accipitrimorphae consistently scores higher, with a more pronounced advantage.

**Table 5.** logL-values for the landbird problem under different conditions.

|  | owl+Accipitrimorphae | owl+Cavitaves |
| --- | --- | --- |
| 12 taxa | -3150.82826 | -3151.45570 |
| 12 taxa(with c) | -2990.72508 | -2991.05644 |
| 5 taxa | -1141.76041 | -1141.89428 |
| 5 taxa(with c) | -1129.37788 | -1129.37355 |
| 5 taxa(with q) | -1458.65812 | -1458.73225 |
| 5 taxa(with q and c) | -1446.93976 | -1447.83584 |
| 33 taxa | -10451.2065 | -10451.3403 |
| 33 taxa(with c) | -9925.93296 | -9926.08197 |

While the phylogenetic placement of owls remains somewhat uncertain, our model favors the owl + Accipitrimorphae clade.

### Waterbird problem: why we need variable *c*?

The phylogenetic relationship of the seven waterbird orders is easy to reconstruct. However, for our main tree of Neoaves on the left of Figure 5 the cherry containing pelican and egret receives an edge length of 0. This is a violation of the assumption of constant insertion rates. Separate trees for the waterbirds with constant and variable introduction rates are shown in Figure 10. A closer examination of the raw data reveals that for egret, pelican and ibis, only two markers support the egret+pelican clade, with one marker each supporting egret+ibis and pelican+ibis. Higher in the tree, there are 227 markers supporting the clade of all waterbirds except for the loon.

**Fig. 10:**
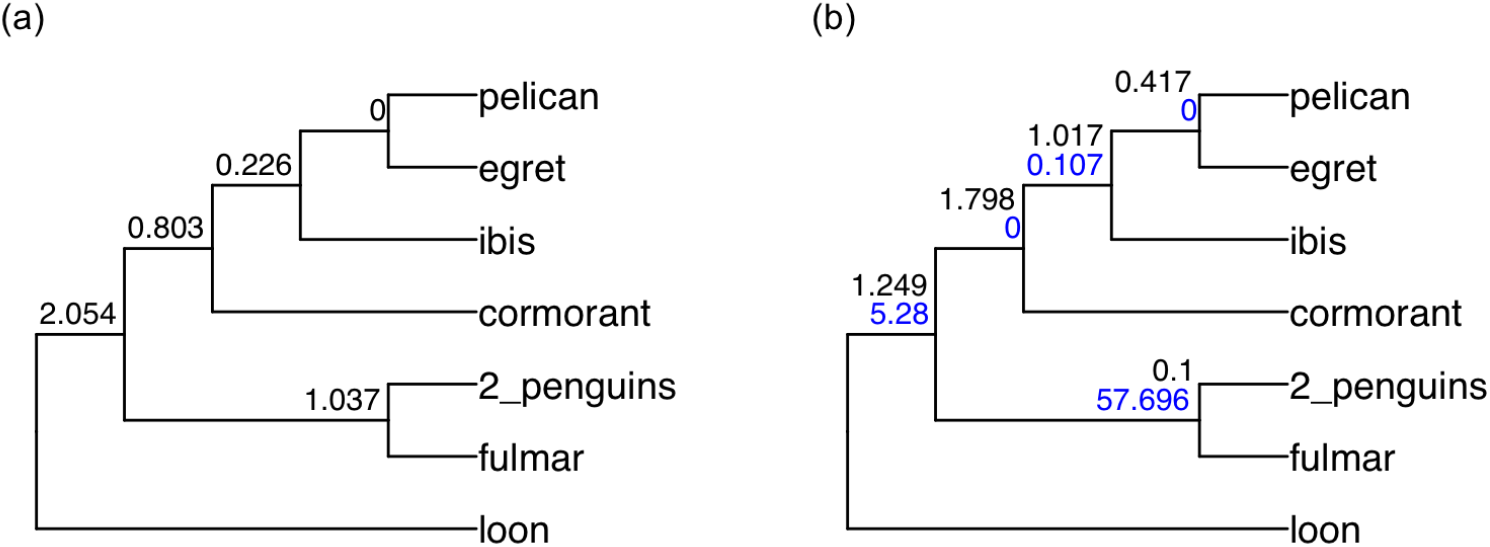
Waterbird Trees. (a) constant *c*; (b) variable *c*.

Consequently, if assuming equal insertion rates across all branches, the egret+pelican branch shows zero length due to insufficient marker support. However, introducing variable insertion rates *c* (Figure 10(b)) causes a branch of length 0.47 with insertion rate zero. Indeed there is a chain of three edges with zero or very low introduction rates and edge lengths longer than 0.4.

The decreasing introduction rate of retrotransposons in the data set was discussed in detail in Suh et al. [2012]. They found that the retroposons causing the long terminal repeats that were discovered from the reference genomes of chicken and zebra finch were very active during the time of the fast Neoavian radiation. This activity had already faded when the lineage leading to ibis, pelican, and egret had branched off other waterbirds.

This example shows that insertion rates may differ for other reason than population dynamics.

## Discussion

This study introduces the MLRE method for phylogenetic tree reconstruction, addressing challenges posed by incomplete lineage sorting (ILS) and marker distribution heterogeneity. The results demonstrate that MLRE achieves higher accuracy compared to traditional methods, particularly when integrating parameter *c*. This parameter plays a crucial role in handling uneven marker distributions, enabling the detection of branches that might otherwise remain unresolved. Moreover, the MLRE method can be used to measure the phylogenetic trees inferred in other studies, providing a unified framework for evaluating tree support under our model.

Compared to previous methods, the new approach requires slightly more computational time than Astral BP but less than SDP Quartets. Most of the time is used for the SPR tree search. For the neoaves data, no SPR move is performed and the first NNI-optimal tree happens to be the final solution. However, we have observed local optima with respect to NNI for other data sets, so the SPR search is recommended. Using the missing data parameter *q* can only be applied to problems with a relatively small number of taxa (below 10). It is useful to check the position of some taxa with an unusual high or low missing data ratio but should not be routinely included.

Whether it is worth to allow variable insertion rates, depends on how much information there is in the data. One way to measure this is the Bayesian information content BIC, and this is frequently used for model selection by maximum likelihood methods like the model finder in IQtree [Kalyaanamoorthy et al., 2017]. The BIC will usually improve when variable insertion rates are allowed. However, in order to avoid overfitting, it is worth to not allow the parameters, if they tend to be extreme (either close to 0 or very large for many edges) for an optimal solution.

It is worth emphasizing that MLRE is not a marker-level adaptation of the multi-species coalescent (MSC) model. We use polymorphic as a state of a marker without considering the percentage of individuals within a population that carry it. Realistically, a marker is introduced in a single individual, so it is much more likely to get lost than to get fixed early on. However, since all markers are observed in some present day species, they must have reached a significant proportion at some point. Assuming that a polymorphic marker is equally likely to get fixed or lost is a justifiable simplification which has also been made by Polymorphism Parsimony [Felsenstein, 1989], MPRE [Suh et al., 2015], and by the marker simulation in Suh [2016]. Nevertheless, the performance drop of MLRE between the data sets simulated here and by Molloy et al. [2022] is probably due to this assumption.

### Which data type should we trust?

There are only 390 markers that are informative between the 13 Buneman clusters of the Neoaves data set, so a few unlucky markers can easily change the tree. Full genomes, on the other hand, with tens of thousands of loci appear to be much more informative. However, the recently published big studies in Wu et al. [2024] and Stiller et al. [2024] only share five out of eleven clades resolving the Buneman clusters. This shows that even with many full genomes wrong trees can be reconstructed. The obvious advantage of retrotransposon data is to have very little (if any) homoplasy. It is debatable how much weight should be given to the results of a retrotransposon data analysis, if it conflicts with large studies of aligned DNA.

For the earliest branches of the Neoaves phylogeny, the MLRE trees have low support and Pteroclimesites branch off alone, instead of forming the Columbimorphae clade together with pigeons that is very stable throughout recent studies. This is an example, where the lack of markers might cause an error in the MLRE trees, even though Figure S2 in Mirarab et al. [2024] shows that an outlier region strongly supports Columbimorphae. The clusters with higher support, like Aequornithes + Eurypqimorphae + Telluraves + Strisores + Charadriiformes + Opisthocomiformes, should be regarded a plausible hypothesis, on par with clades suggested by whole genome analyses. The MLRE scores of published trees from other data, as shown in Table 4, quantifies the disagreement between those trees and our model, suggesting that the trees by Jarvis et al. [2014] and Stiller et al. [2024] are much more plausible than the ones by Prum et al. [2015] and Wu et al. [2024]. Finally, when the retrotransposon data is as clear as for the mousebird position, it should overrule other data types, unless homoplasy turns out to be not as rare as currently believed.

### Neoaves phylogeny

Even though the title of Jarvis et al. [2014] indicated otherwise, the Neoaves phylogeny was not resolved in 2014, and it has not been since. Nevertheless, there has been significant progress from the newly available data. The Buneman clusters of the Suh et al. [2015] retrotransposon data partitions the 36 orders of modern birds into 13 non-controversial clades. Reddy et al. [2017] used a consensus tree to group Pteroclimesites and pigeons as well as Musophagiformes, Otidiformes, and Cuculiformes together into Columbimorphae and Otidimorphae, respectively, and called the remaining 10 groups the “magnificent seven” and three “orphaned orders”. Columbimorphae have been recovered by most recent studies, but not by Suh et al. [2015], Kuhl et al. [2020], or by our MLRE trees. Otidimorphae are also found frequently, but not by Suh et al. [2015], Reddy et al. [2017], Kuhl et al. [2020], or our MLRE trees with fixed insertion rate. The resolution of the ten remaining major clades is still controversial, and Suh [2016] even suggested a 9-way polytomy which only contains another frequently observed cluster, the Phaethoquornithes. Our analysis indicates that there is resolution, and there is significant support that Pteroclimesites, Columbiformes, Mirandonithes, and Otidimorphae branched off early. While the exact order of that process is unclear and the position of Gruiformes varies, the remaining Neoaves form a well supported clade. These tendencies are shared by Jarvis et al. [2014] and Stiller et al. [2024], which explains the relatively good score of their trees in Table 4. The resolution of our proposed cluster containing Aequornithes, Eurypqimorphae, Telluraves, Strisores, Charadriiformes, and Opisthocomiformes is too unstable to confidently hypothesise any groupings.

Within Telluvares (landbirds), the mainstream hypothesis is that they first split into Australaves and Afroaves (exceptions are Prum et al. [2015], Houde et al. [2019], and Wu et al. [2024]), and that is confirmed by MLRE. Then our method very confidently divides Afroaves into mousebirds and a cluster of owls, Accipitrimorphae, and Cavitaves. As mentioned earlier this split is rarely found by other data types, for example the UCE tree in Jarvis et al. [2014]. Resolving the latter group is the most interesting open question of landbird phylogenetics. Our method, as well as Stiller et al. [2024], slightly prefer grouping together owls and Accipitrimorphae. There is significant support for a group of owls with Cavitaves, but since the third possible grouping is unsupported, this does not indicate a trifurcation (as suggested by Suh et al. [2015]) but a reticulation.

Within Otidimorphae, our method always finds a cluster of Otidiformes and Musophagiformes which has been named Musophagotides [Sangster et al., 2022]. This clade is stronger than Otidimorphae, but it is not found by Stiller et al. [2024].

### Future work

While major sequencing efforts, most notably the Bird 10K Genomes Project [Zhang et al., 2015], have greatly expanded the availability of avian genomic data, retrotransposon marker datasets have not been updated. In Feng et al. [2020], a data set of 363 bird genomes was published, which was used for the phylogenetic analysis in Stiller et al. [2024]. Following the pipeline of Suh et al. [2015] and then making consensus sequences for all orders, has the potential to reduce the percentage of missing data. The data set analysed here contains two eagles with 162 and 135 question marks, respectively, and their consensus sequence contains only 105. This shows that even closely related species can add a lot of information to a reteotransposon data set. In addition, better taxon sampling might increase the number of markers, because candidate loci that were filtered out for being phylogenetically uninformative are now found in more than one order. In addition, there is progress on generating retroelement insertion data sets [Churakov et al., 2020], and MLRE could be applied to other types of low-homplasy markers like large indels [Schull et al., 2022].

As mentioned before the support for conflicting positions of owls within the landbirds tree indicates a reticulation. Indeed, whenever for three taxa, the second best rooted triple has significantly more support than the worst one, this cannot be explained by ILS alone. In Springer et al. [2020], an asymmetry test is applied to retroelement data to predict gene flow. Using the model of MLRE to score phylogenetic networks and trying to find a network with optimal score is a challenging task.

Finally, it would be desirable to modify MLRE such that it becomes consistent under the multi-species coalescent (MSC) model. This would require that the probability of the loss of a marker initially is much larger than the probability of fixation. As long as a marker stays polymorphic, the ratio between the probabilities needs to gradually increase, converging to 1. This means that the transition probability matrix for a fixed edge is not constant, thus it seems hard to develop an algorithm following this approach.

## Supplementary Material

The codes and datasets of this study have been uploaded to [https://github.com/gaoliangs/MLRE] for public access and use.

