## Supplementary Material for "A Maximum-Likelihood Method to Construct Phylogenetic Trees Using Low-Homoplasy Markers: New Insights into Neoaves Phylogeny"

This supplementary material contains a series of tables and figures that support and extend the findings presented in the main text.

- Table S1 summarizes the additional number of markers required to reconstruct specific clades, along with the bird taxa included in each corresponding cluster.
- Figure S1 shows the true tree used in the simulation experiments, serving as a reference for evaluating the performance of different phylogenetic reconstruction methods.
- Figure S2 uses the dataset from [5] and compares the reconstruction results obtained by two different methods.
- Figure S3 presents simulation results designed to test the robustness of our approach under controlled conditions.
- Figures S4–S11 display branch lengths inferred by MLRE on several previously published Neoaves phylogenies, allowing direct comparison with existing topologies.
- Figure S12 shows an ASTRAL_BP tree with outgroup taxa included, highlighting the impact of rooting on tree topology.
- Figure S13 illustrates branch lengths inferred by MLRE on the tree constructed using the Dolpenny method [1].
- Figure S14 represents the number of congruent markers supporting each branch of the MLRE tree.
- Figure S15 shows a simplified Neoaves tree containing only 13 taxa.

Together, these figures offer additional validation, comparison, and context for the phylogenetic inferences discussed in the main manuscript.

**Table S1.** Number of markers added

| Cluster | Taxa in Cluster | flexible c | constant c |
|---|---|---|---|
| Otidae [3] | Otidimorphae+Strisores | 3 | 3 |
| Otidimorphae [3] | cuckoos, bustards, turacos | 0 | 1 |
| Columbea [3] | Columbimorphae+Mirandornithes | 8 | 4 |
| Columbimorphae [3] | pigeons, mesites, sandgrouse | 2 | 2 |
| Gruae [3] | Opisthocomiformes+Gruimorphae | 3 | 4 |
| Phaethoquornithes [3] | Eurypygimorphae+Aequornithes | 1 | 1 |
| Inopinaves [6] | Opisthocomiformes+Telluraves | 6 | 3 |
| Gruimorphae [8] | Gruiformes+Charadriiformes | 3 | 3 |
| Columbaves [6] | Columbimorphae+Otidimorphae | 9 | 9 |
| Litusilvanae [10] | Strisores+Gruimorphae | 3 | 5 |
| Elementaves [8] | Gruae+Strisores+Phaethoquornithes | 4 | >10 |
| Aequorlitornithes [10] | Mirandornithes+Opisthocomiformes+Phaethoquornithes | >10 | >10 |

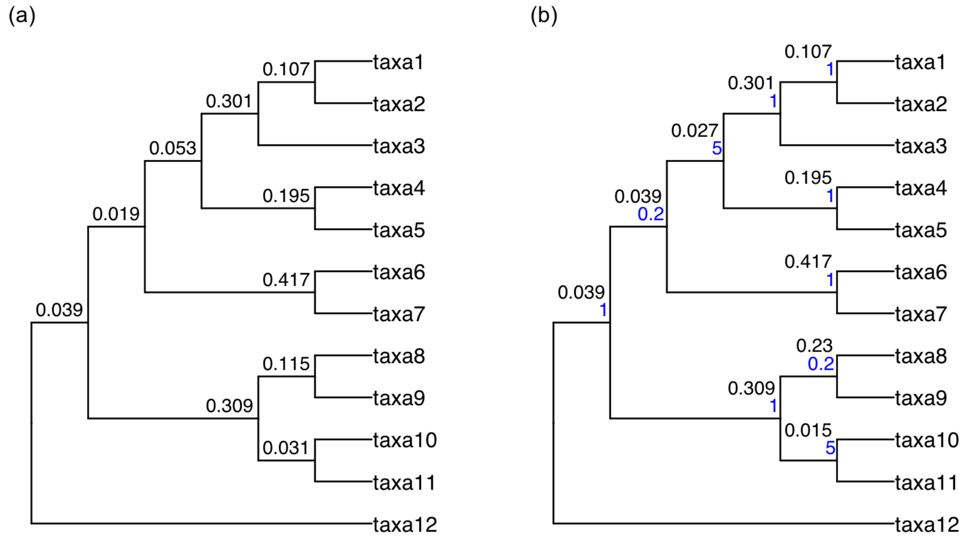


**Fig. S1. Simulated tree from adjusted palaeognathae.** (a) The tree used for simulations under the original model, (b) The tree used for simulations with variable insertion rates. Black numbers indicate the branch lengthes, and blue numbers indicate insertion rates. The phylogenetic trees in this supplementary material are all plotted using the R package ggtree [11].

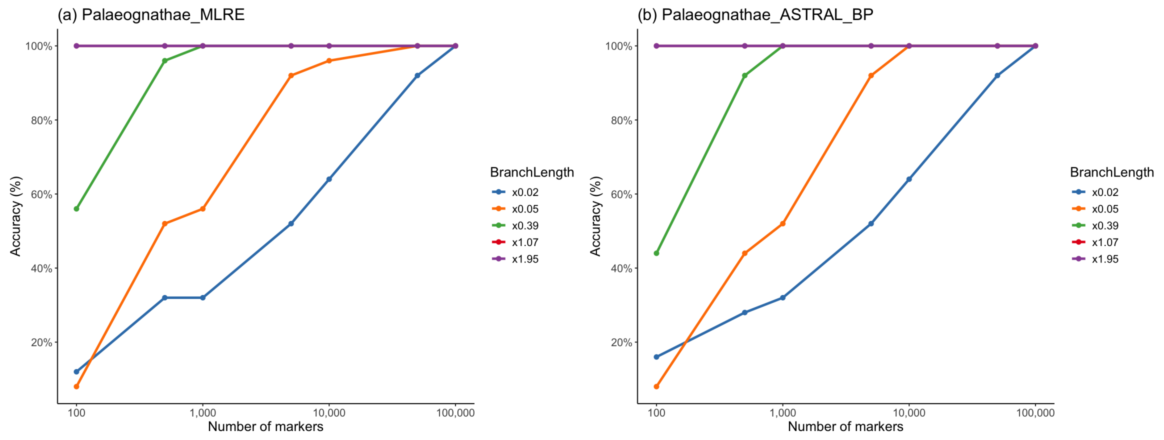


**Fig. S2.** Performance comparison on simulated data [5] generated by ms. (a) Results from the MLRE method, (b) Results by ASTRAL_BP. Different colors represent different branch lengths. The x-axis indicates the data size, and the y-axis represents the precentage of correctly recovered branches.

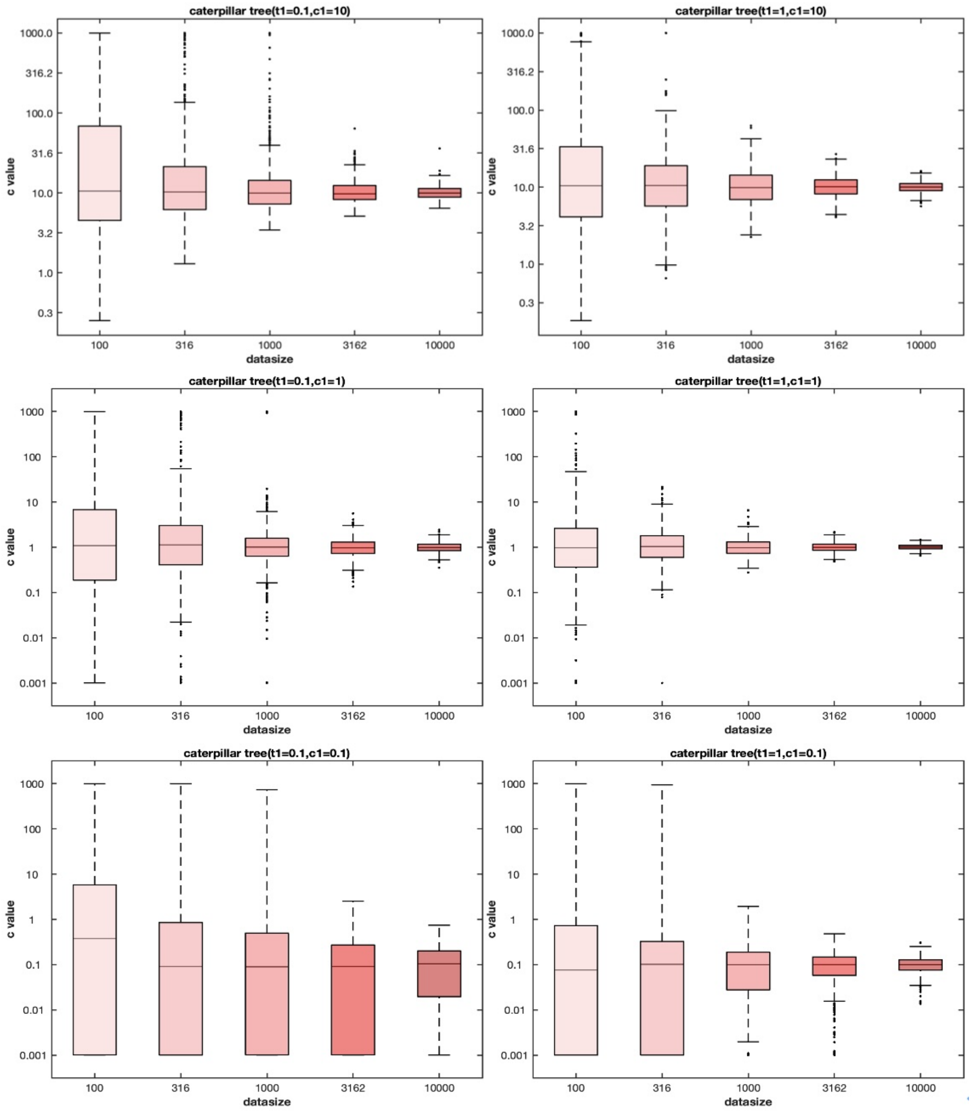


**Fig. S3. Distribution of estimates of insertion rate $c$ for different data sizes.** Left and right represent simulated trees with branch lengths of 0.1 and 1, respectively. Top, middle, and bottom represent simulated values of 10, 1, and 0.1, respectively, for the insertion rate $c$. The x-axis is the number of markers ranging from 100, 316, 1,000, and 3,162 to 10,000; and the y-axis is the distribution of the estimated value of the insertion rate $c$, which is constrained to the range between (0.001, 1,000). Note that the y-axis is scaled differently.

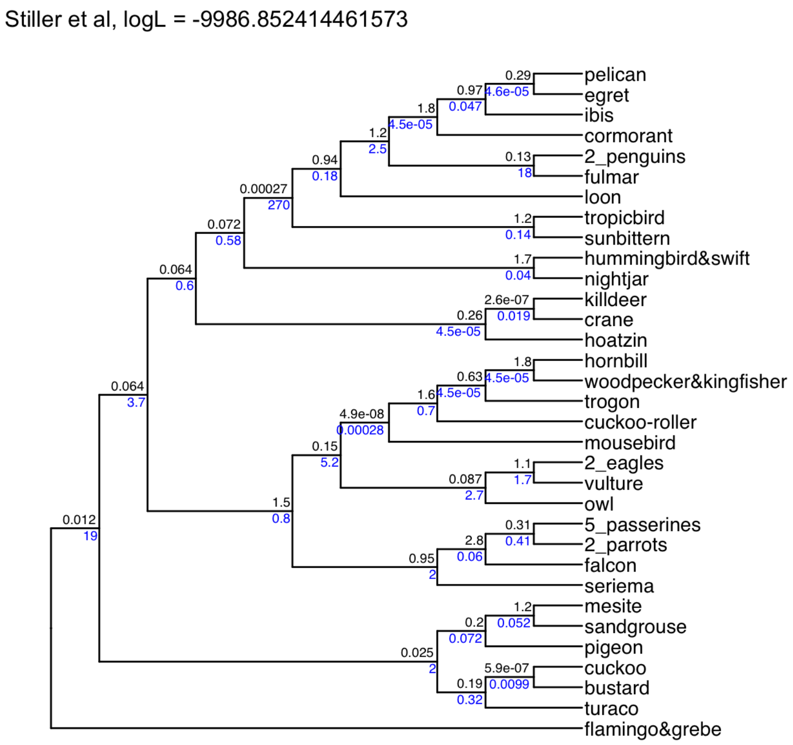


**Fig. S4.** The tree topology is from [8], with branch lengths inferred using the MLRE method.

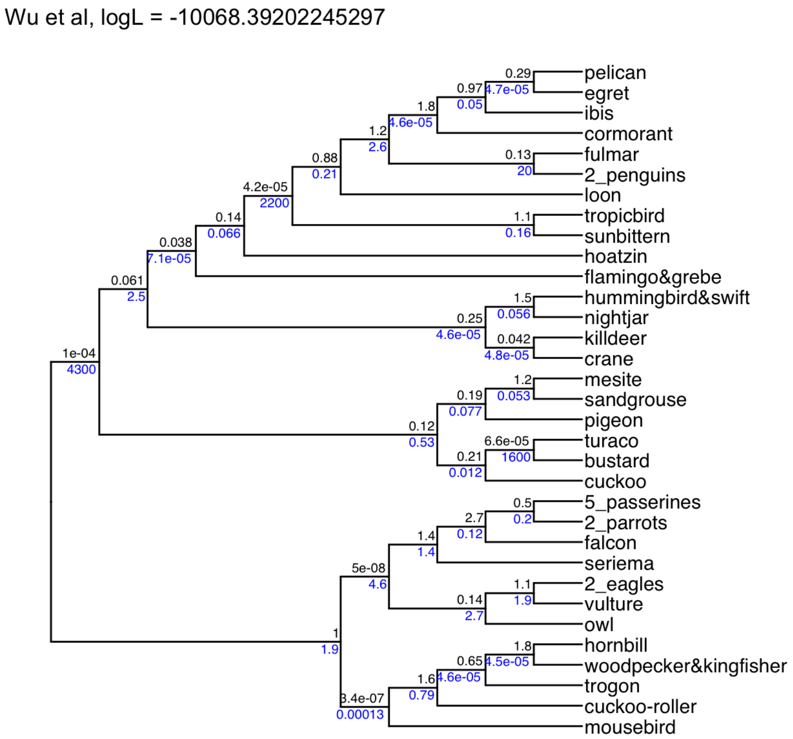


**Fig. S5.** The tree topology is from [10], with branch lengths inferred using the MLRE method.

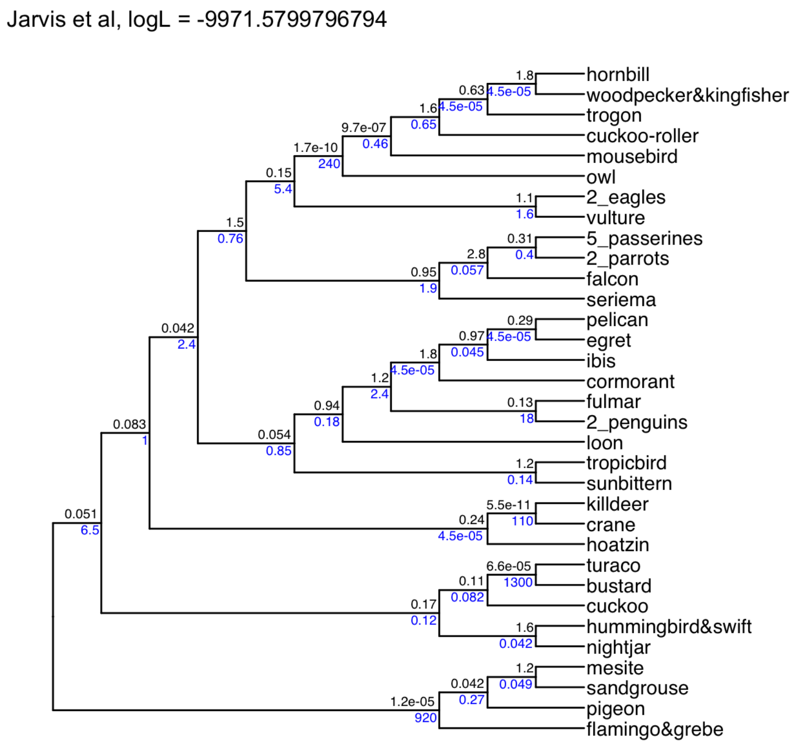


**Fig. S6.** The tree topology is from [3], with branch lengths inferred using the MLRE method.

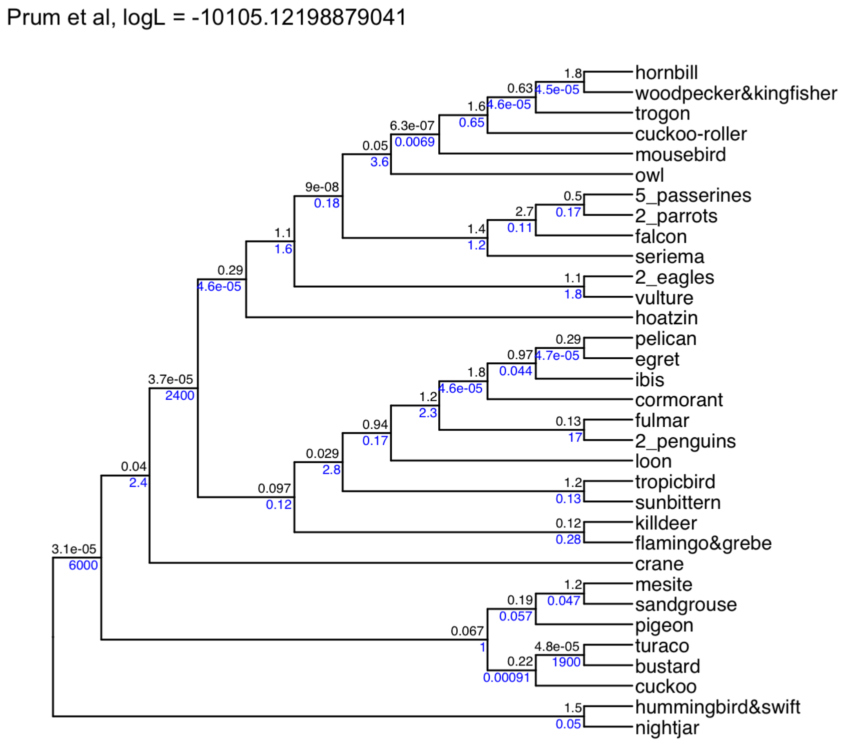


**Fig. S7.** The tree topology is from [6], with branch lengths inferred using the MLRE method.

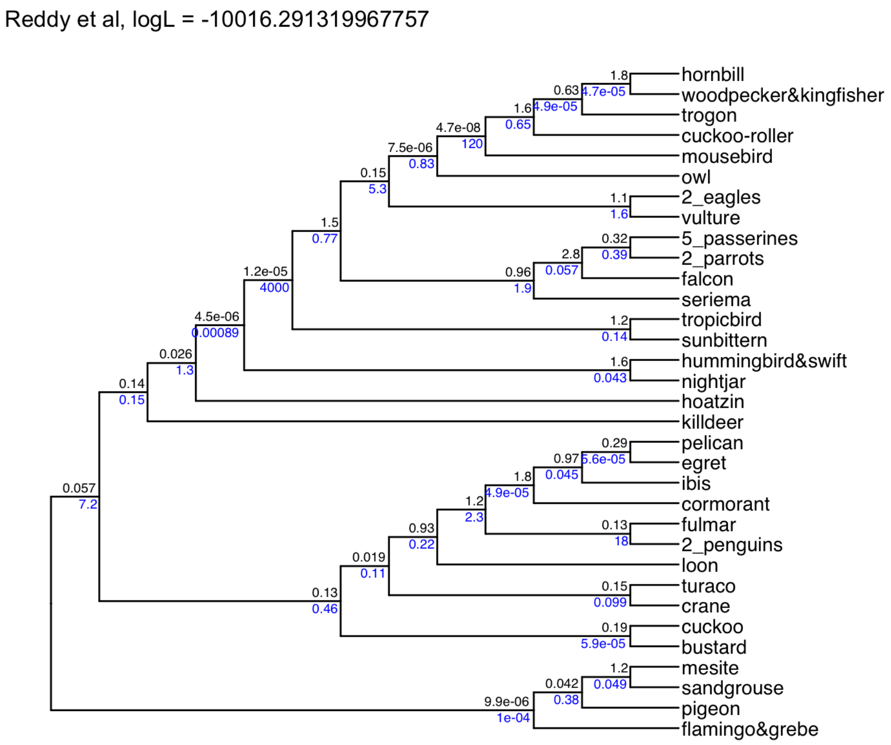


**Fig. S8.** The tree topology is from [7], with branch lengths inferred using the MLRE method.

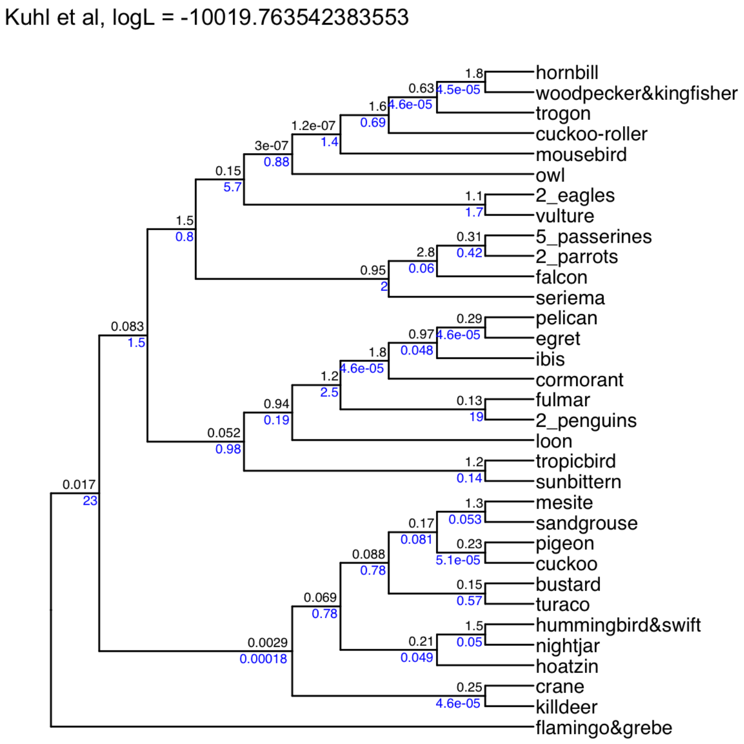


**Fig. S9.** The tree topology is from [4], with branch lengths inferred using the MLRE method.

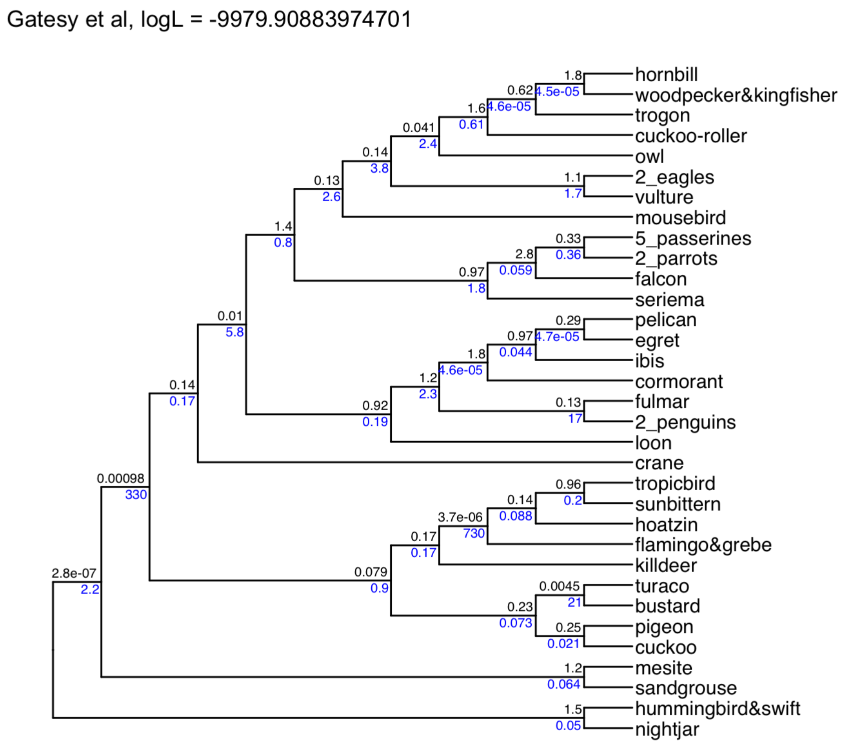


**Fig. S10.** The tree topology is from [2], with branch lengths inferred using the MLRE method.

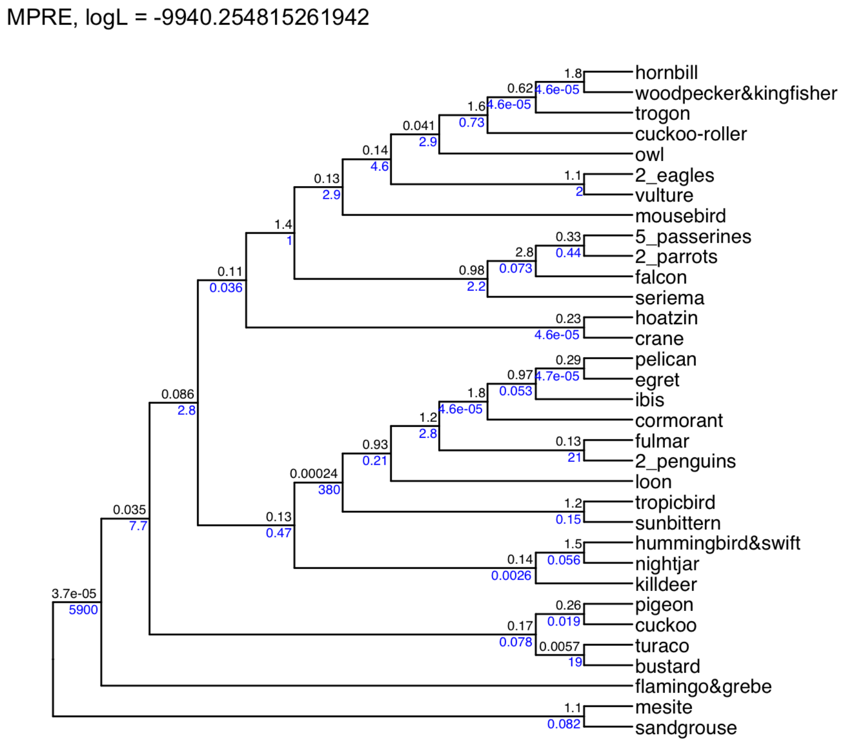


**Fig. S11.** The tree topology is from [9], with branch lengths inferred using the MLRE method.

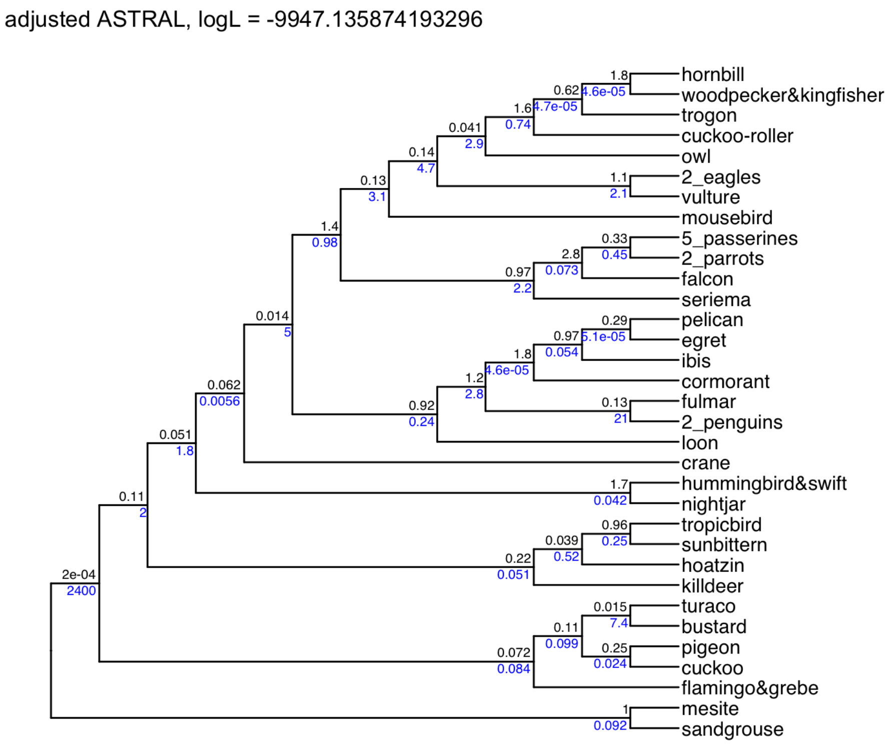


**Fig. S12. ASTRAL_BP adjusted.** To root the tree, an all-zero outgroup was added to the dataset of 48 Neoaves taxa, and the phylogeny was inferred using the ASTRAL_BP method.

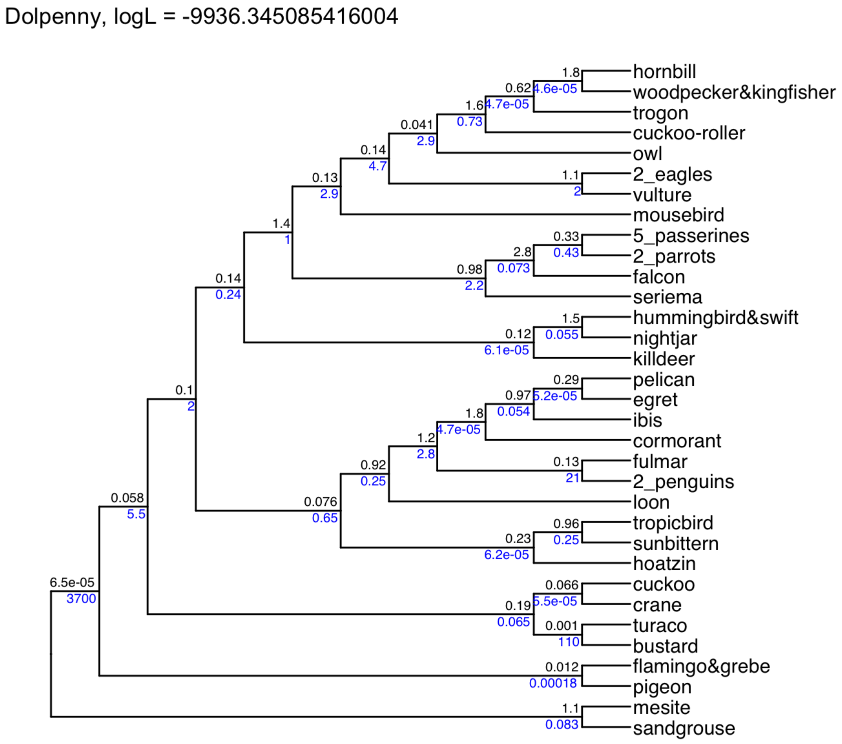


**Fig. S13.** The tree is generated from dolpenny method [1], with branch lengths inferred using the MLRE method.

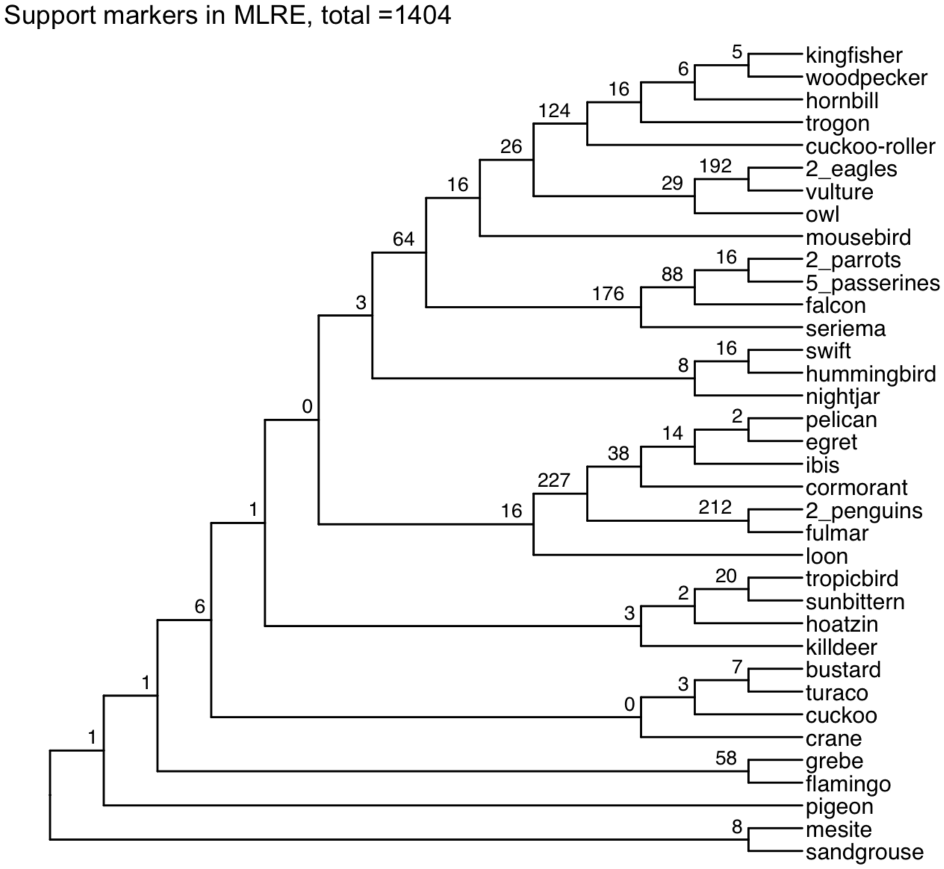


**Fig. S14. Congruent markers with the MLRE tree.** For each branch of the MLRE tree, we calculated the number of congruent markers. There are 1404 such markers compare to 1398 markers for MPRE method, MLRE consistently results in a higher number of congruent markers.

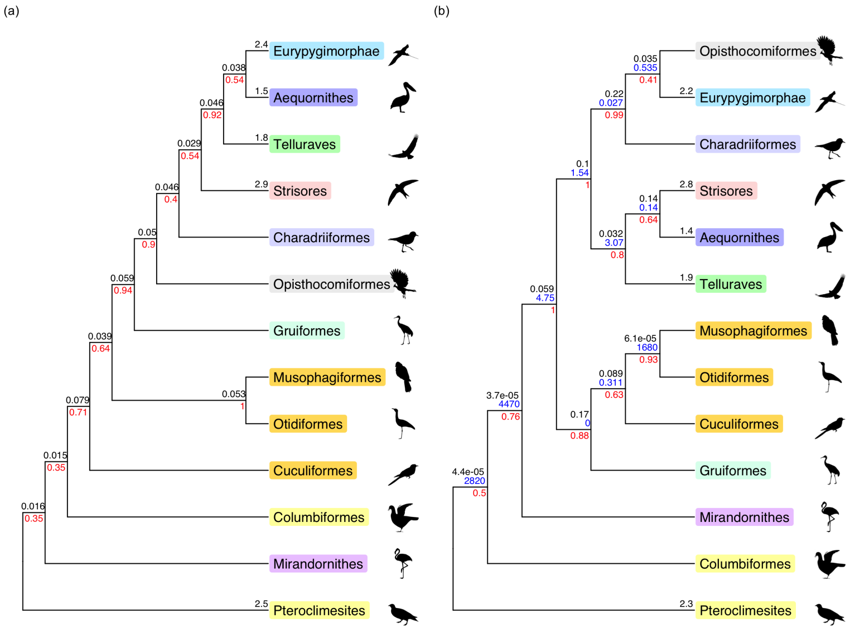


**Fig. S15. MLRE tree for 13 Neoaves taxa.** (a) The Neoaves tree under the MLRE(c=1), logL = -3142.83593; (b) The Neoaves tree with variable insertion rates, logL = -3114.30806. Black numbers indicate branch length, blue numbers indicate insertion rates, and red numbers indicate branch support. In particular, only the evolutionary branches corresponding to the group-taxa have finite leaf edge length.
